# Predicting cell population-specific gene expression from genomic sequence

**DOI:** 10.1101/2022.09.06.506837

**Authors:** Lieke Michielsen, Marcel J.T. Reinders, Ahmed Mahfouz

## Abstract

Most regulatory elements, especially enhancer sequences, are cell population-specific. One could even argue that a distinct set of regulatory elements is what defines a cell population. However, discovering which non-coding regions of the DNA are essential in which context, and as a result, which genes are expressed, is a difficult task. Some computational models tackle this problem by predicting gene expression directly from the genomic sequence. These models are currently limited to predicting bulk measurements and mainly make tissue-specific predictions. Here, we present a model that leverages single-cell RNA-sequencing data to predict gene expression. We show that cell population-specific models outperform tissue-specific models, especially when the expression profile of a cell population and the corresponding tissue are dissimilar. Further, we show that our model can prioritize GWAS variants and learn motifs of transcription factor binding sites. We envision that our model can be useful for delineating cell population-specific regulatory elements.

## INTRODUCTION

In multicellular organisms, every cell has the same DNA apart from somatic mutations. Yet its function and the related proteins and genes expressed vary enormously. This is among others caused by transcriptional and epigenetic regulation. Proteins that bind the DNA sequence around the transcription start site (TSS) control whether a gene is transcribed in a cell (1, 2). Which transcription factors, and thus which DNA binding motifs, are essential differ per cell population (1–4). As such, mutations in regulatory regions might affect specific tissues or cell populations differently. Improving our understanding of these regulatory mechanisms will help us relate genomic functions to a phenotype.

For example, while promoter sequences are identical across the four major human brain cell populations (neurons, oligodendrocytes, astrocytes, and microglia), almost all enhancer sequences, the regions in the DNA where a transcription factor binds, are cell population-specific (3). These population-specific regulatory elements are discovered by combining single-cell measurements of different data types, including chromatin accessibility, ChIP-seq, and DNA methylation. Bakken et al., for instance, identified differentially methylated and differentially accessible regions across neuronal cell populations in the human brain, albeit with little overlap (5). This emphasizes the complexity of transcriptional regulation and the need for more measurements to fully resolve these mechanisms at the cell population-specific level.

An alternative approach would be to train a computational model that directly predicts gene expression from the genomic sequence around the TSS. This way, we can learn which regulatory elements are important for transcriptional regulation in different contexts. Several computational methods have been developed for this task (6–12). These methods have in common that they one-hot encode the DNA sequence and input this to either a convolutional neural network (CNN) or transformer. ExPecto, Xpresso, and ExpResNet predict expression measurements from bulk RNA-sequencing, while Basset, Basenji, BPNet, and the Enformer model predict regulatory signals, such as cap analysis gene expression (CAGE) reads or TF binding from CHIP-nexus.

A promising application of these models is to prioritize variants that have been identified using genome-wide association studies (GWAS) (6, 13). Using GWAS many potential disease-associating variants have been identified (14–16). Within each locus, however, it is often challenging to pinpoint which variant is causal and which gene is affected by the variant.

These current computational gene prediction models, however, are designed for predicting bulk gene expression data. This means that they are either tissue-specific or could be applied to FACS-sorted cells (13). Since transcriptional regulation is even more context-specific, the resolution of current methods is not sufficient for heterogeneous tissues where single-cell RNA-sequencing (scRNA-seq) has revealed hundreds of cell populations (5, 17, 18). To increase the resolution, the models would ideally be trained on scRNA-seq data.

Here, we present scXpresso, a deep learning model that uses a CNN to learn cell population-specific expression in scRNA-seq data from genomic sequences. Since single-cell and bulk data have different characteristics and distributions, we explored whether this type of model is suitable for single-cell data. We show that (i) cell population-specific models outperform tissue-specific models on several tissues from the Tabula Muris, (ii) increasing the resolution improves the predictions for human brain cell populations, and (iii) in-silico mutagenesis of the input sequence can be used to prioritize GWAS variants.

## MATERIAL AND METHODS

### Architecture of scXpresso

scXpresso is a one-dimensional convolutional neural network (CNN) adapted from the (bulk gene expression-based) Xpresso model (9) (Figure 1A, S1). Compared to Xpresso, we designed scXpresso as a multitask model so that it predicts the expression of multiple cell populations simultaneously. The input to the CNN is four channels with the one-hot encoded sequence around the transcription start site (TSS) (7kb upstream and 3.5kb downstream). Every channel represents one of the four nucleotides (A, C, T, G). For some positions, the exact nucleotide is not known (e.g. any nucleic acid (N) or a purine nucleotide (R)). The exact coding scheme for such positions is shown in Table S1. The CNN consists of two convolutional layers. The output of the convolutional layers is flattened and concatenated with the half-life time features. Different from Xpresso, we decreased the number of half-life time features from eight to five; the three features we removed (5’ UTR, ORF, and 3’ UTR GC content) correlated less with half-life time, so we removed them to make the model less complex (9, 19, 20). Together, this is subsequently fed into a fully connected (FC) layer. For the multitask model, there is only one FC layer. For the other models, which we use to make tissue-specific predictions as a comparison, we used two FC layers. The output of the FC layers is the aggregated expression per tissue or for each cell population.

**Figure 1.**
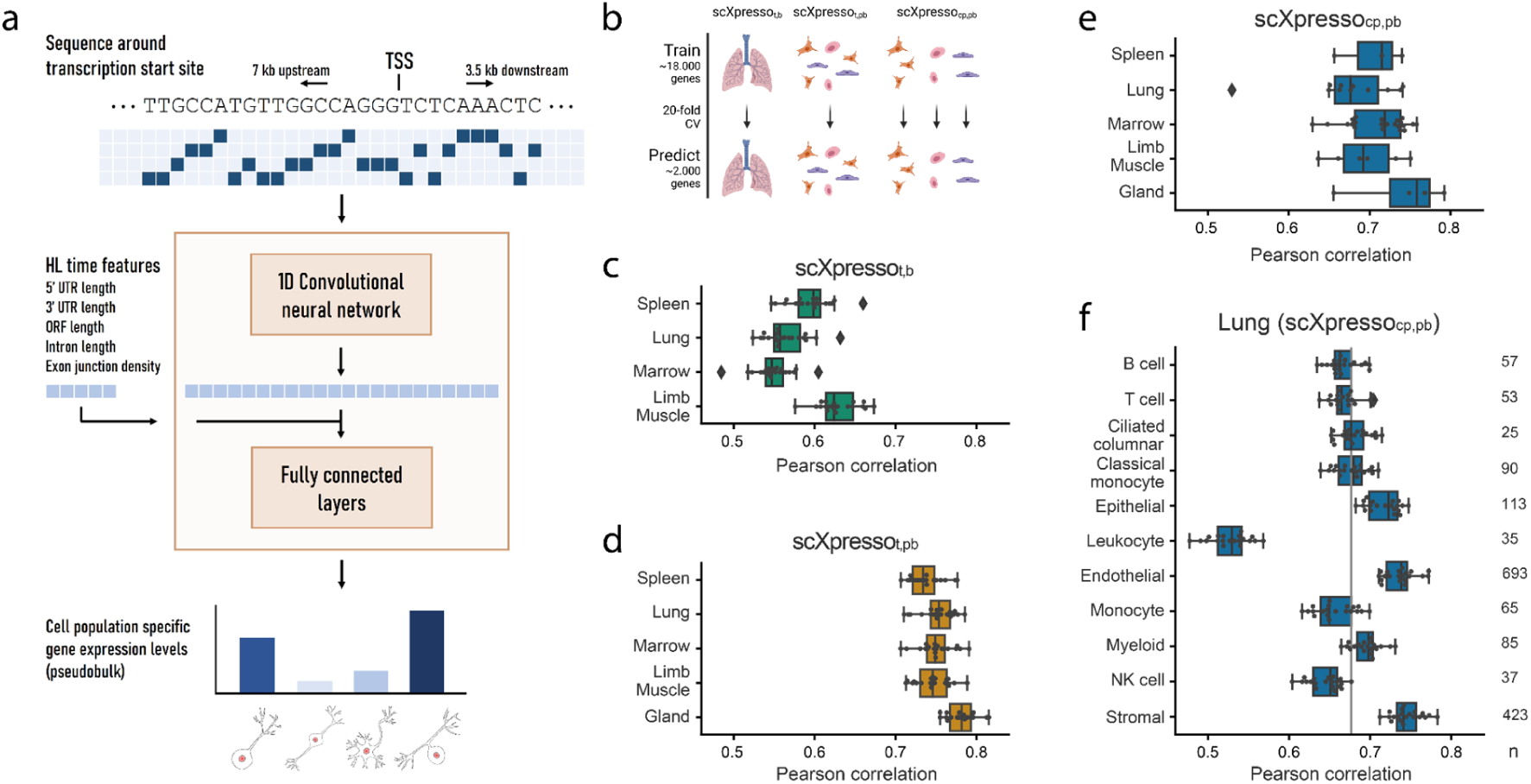
Schematic overview of scXpresso and performance on Tabula Muris datasets. **a)** We one-hot encode the DNA sequence around the transcription start site (TSS) and input this to a one-dimensional convolutional neural network (CNN). The output of the CNN is flattened and concatenated with the five half-life time features. The fully connected layers output the cell population’s specific gene expression levels simultaneously (Figure S1, see **Methods**). **b)** Schematic overview of the experiment. **c-d)** Performance of scXpresso_t, b (_tissue-specific (t) model on bulk (b) data) and scXpresso _(_tissue-specific model on pseudo-bulk (pb) data), respectively. Every dot is the performance (Pearson correlation) across one fold of the 20-fold CV. **e)** Performance of scXpresso_cp, pb (_cell population-specific (cp) model on pseudo-bulk data) summarized per tissue. Every dot represents the model’s performance on a cell population in that tissue (median Pearson correlation across the 20 folds). **f)** Performance of scXpresso_cp, pb o_n the different lung cell populations. The grey line indicates the median performance across all cell populations. Every dot is the performance across one fold of the 20-fold CV.

### Training scXpresso

We split the genes into a train, validation, and test dataset and do 20-fold cross-validation. These sets are the same across all experiments (i.e. one train, validation, and test set for mouse genes and one for human genes) such that the results of different models can be compared. We update the weights of scXpresso using the Adam optimizer based on the mean square error loss on the training set. The initial learning rate is set to 0.0005 and if the loss on the validation set is not improved from 5 epochs, the learning rate is reduced by a factor of 10. We train the model for 40 epochs and the model with the lowest loss on the validation set is used for evaluation on the test dataset. Since there is always some stochasticity when training a CNN, we always train 5 models and average the predictions. We used the following software packages for training the model: Pytorch (version 1.9.0) (21), CUDA (version 11.1), cuDNN (version 8.0.5.39), and Python (version 3.6.8).

### Datasets

#### Tabula Muris

The single-cell Tabula Muris data (22) for the five different tissues (gland, spleen, lung, limb muscle, and bone marrow) and two different protocols (10X and FACS-sorted Smart-seq2) were downloaded from: https://figshare.com/projects/Tabula_Muris_Transcriptomic_characterization_of_20_organs_and_tissues_from_Mus_musculus_at_single_cell_resolution/27733. To extract input features, we downloaded the reference genome (MM10-PLUS) that was used during the alignment from: https://s3.console.aws.amazon.com/s3/object/czb-tabula-muris-senis?region=us-west-2&prefix=reference-genome/MM10-PLUS.tgz.

The four bulk datasets (spleen, lung, limb muscle, and bone marrow) from the Tabula Muris were downloaded from https://www.ncbi.nlm.nih.gov/geo/query/acc.cgi?acc=GSE132040 (23). For the bulk data, we used the same reference genome as for the single-cell data.

#### Human motor cortex data

The human motor cortex data from the Allen Institute (5) was downloaded from the Cytosplore Comparison Viewer. We downloaded the reference genome (version GRCh38.p2) and corresponding GTF file with information about the location of transcription start sites of the genes here: (https://www.gencodegenes.org/human/release_22.html).

### Aggregated expression values

First, we normalized the count matrices. For the single-cell datasets, we did library size normalization in the same way as The Tabula Muris Consortium: i.e. counts per million for the smart-seq2 data and counts per ten thousand for the 10X data (22). For the bulk Tabula Muris data, we performed TPM normalization. For the single-cell datasets, we used the annotations defined by the authors to aggregate the expression values per tissue or per cell population using log_10_ *mean(x) (without pseudocount)* into pseudobulk values. The advantage of not adding a pseudocount is that the distribution looks more like a normal distribution, which makes it easier to train the models (Figure S2). A limitation, however, is that we could not calculate the exact value for genes that were not expressed in any of the cells. For these genes, we replaced the pseudobulk values with -4 in the Tabula Muris and -5 in the motor cortex dataset, since this extrapolated well (Figure S2). For the bulk data, we aggregated over the samples instead of the cells. Here, we set the genes that are not expressed in any of the samples to -4. We standardized the expression values before running the model such that the average expression of all genes in each cell population or tissue is zero and the standard deviation is one. Before analyzing the results and comparing the predictions across cell populations, we undid the z-score normalization but kept the log normalization.

### Input features

#### Sequence around the transcription start site

Before extracting the sequences around the transcription start site, we removed genes that are transgenes, ERCC spike-ins, genes without a coding region, and genes on the Y chromosome. This resulted in 20, 467 mouse genes and 18, 138 human genes. Some genes had multiple transcripts. We downloaded a list with canonical transcripts for each gene from biomart and we used the transcript and transcription start site belonging to the canonical transcript. If the canonical transcript was not defined, we used the transcript that had the longest coding region. After having defined the transcription start site for each gene, we used seqkit (24) to extract sequences from the FASTA file containing the reference genome.

#### Half-life time features

For every gene, we extracted five half-life time features: 5’ UTR length, 3’ UTR length, ORF length, intron length, and exon junction density 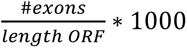. We obtained these features by first filtering the GTF files for the canonical or longest transcript. The 5’ UTR length is the length of the sequence from the start of the first exon to the start codon. The 3’ UTR length is the length of the sequence from the last coding sequence to the end of the last exon. The ORF length is the sum of the length of the coding sequences. The intron length is the length of the transcript minus the length of the ORF, 5’ UTR, and 3’ UTR. All features are log-normalized using log_10_*(x* + 0.1) and afterwards z-scaled.

### Evaluating the predictions

For every gene in the test dataset, we averaged the predictions of the five models we trained. We evaluated the performance for every cell population by calculating the Pearson correlation between the true and predicted expression of the genes in the test set. To evaluate the increase in performance between the tissue-specific (t) pseudobulk (pb) and cell population-specific (cp) pseudobulk (pb) model on the Tabula Muris datasets, we calculate: *Δ_cp_*_, *t*_ *= median Pearson correlation (scEP_cp_*_, *pb*_*) – median Pearson correlation (scEP_t_*_, *pb*_). On the motor cortex dataset, we also evaluated the performance of each gene by calculating the Pearson correlation between the true and predicted expression per cell population.

### In-silico saturation mutagenesis

For *CACNA1I*, we mutated all positions *in-silico*, which means we tested all possible substitutions at every position. We undid the z-score normalization and calculated the difference between the original (wild-type) prediction and the mutated prediction. The prediction models used during these experiments were the models where *CACNA1I* itself was originally in the test set. For every position, we only plotted one predicted difference in expression in Figure 4E. This is the substitution that was predicted to have the largest absolute effect. We downloaded the locations of the candidate cis-regulatory elements that fall within the input region for *CACNA1I* from screen registry v3 (release date 2021) (25). When plotting the difference between two cell populations, we ignored the positions where one is positive and the other predicts a negative effect. This rarely happened and if it was the case, the predicted effect was very small.

### Comparison to other models

#### Enformer

Enformer uses the DNA sequence to predict reads for 5, 313 human tracks which include CAGE, DNAse, CHIP, and ATAC-seq (11). Here, we only looked at the effect of a variant on the CAGE tracks that are related to the brain (77 tracks in total, see Table S2). Enformer predicts the effect of variants on 128bp bins. When predicting the effect of a variant on the CAGE reads, we looked at the effect on the bin containing the TSS.

#### ExPecto

ExPecto predicts gene expression for 218 tissues and cell lines (8). Here, we only focused on 27 outputs that are related to the brain (Table S3). We used the ExPecto web server to predict the effect of the variants (https://hb.flatironinstitute.org/expecto/?tabId=3). ExPecto is trained using Hg19 instead of Hg39. We used the R-package SNPlocs.Hsapiens.dbSNP155.GRCh37 (v 0.99.23) to lift-over the variants. Using ExPecto we could not predict the effect of all variants, since for some variants there was no location in Hg19 found, some were too far away from a TSS, and some were linked to a different gene than we were interested in (see Table S4 for an explanation per variant).

#### Xpresso

We trained the Xpresso model on bulk RNA-seq data from the precentral gyrus (9). The data from two individuals were downloaded from the Allen Human Brain Atlas: https://human.brain-map.org/static/download (H0351.2001, H0351.2002). We used the normalized matrices. Labels were created as described in the Xpresso paper: we took the median expression across the 6 precentral gyrus samples, log-normalized the output using log_10_(*x* + 0.1), and z-score normalized the expression. Similar to scXpresso, we trained the model using 20-fold cross-validation. Per fold, we trained 10 runs and used the model with the lowest MSE on the validation data (as described in (9)). Afterwards, we predicted the effect of the variants. We could not predict the effect of all variants, since some genes were not measured in the bulk RNA-seq data and for some genes, there were no Xpresso input features defined (see Table S4 for an explanation per variant).

## RESULTS

### Predicting cell population-specific gene expression using scXpresso

Here, we present scXpresso, a multitask convolutional neural network (CNN) to predict cell population-specific gene expression using genomic sequences only (Figure 1A, S1). We developed scXpresso by adapting the Xpresso model (9), which was originally designed for bulk data, to single-cell data. Similar to Xpresso, we use two types of input to the model: (1) the DNA sequence around the transcription start site (TSS) (7kb upstream - 3.5kb downstream) to model transcription, and (2) five half-life time features (5’ UTR length, 3’ UTR length, ORF length, intron length, and exon junction density) to model mRNA degradation. We input the one-hot encoded DNA sequence into a CNN. The output of the CNN is concatenated with the half-life time features and fed to a fully connected network (see **Methods**). Since our model is a multitask CNN, the desired output of the fully connected network is the gene expression for every cell population. To obtain one expression value per cell population, we aggregate the single-cell expression into pseudobulk measurements (see **Methods**).

Since single-cell and bulk data have different characteristics, we tested whether scXpresso performs equally well on single-cell and bulk data. We used scRNA-seq data from five different tissues (limb muscle, spleen, gland, marrow, and lung) from the Tabula Muris [13] (Table S5). Here, we used the FACS-sorted cells that were sequenced using the Smart-seq2 protocol. Using the annotations defined by the authors, we aggregate the values per cell population and per tissue into pseudobulk values. For four tissues (limb muscle, spleen, marrow, and lung), there are also bulk RNA-sequencing datasets available (Table S6). We compared the pseudobulk to the bulk expression per tissue and noticed that these are indeed correlated (r_muscle =_ 0.69, r_spleen =_ 0.71, r_marrow =_ 0.50, r_lung =_ 0.67) (Figure S3).

Next, we trained three different models: 1) a tissue-specific (t) model on the bulk (b) values (scXpresso_t, b)_, 2) a tissue-specific model on the pseudobulk (pb) values (scXpresso_t, pb)_, 3) a cell population-specific (cp) model on the pseudobulk values (scXpresso_cp, pb)_ (Figure 1B). The cell population-specific model is, in contrast to the tissue-specific models, a multitask model that predicts the expression of all cell populations in a tissue simultaneously. We evaluated the performance of the models by calculating the Pearson correlation between the true and predicted expression values. In general, the tissue-specific models trained on pseudobulk reach higher performance than the models trained on bulk (Figure 1C-D). Even though the bulk and pseudobulk values are correlated, the pseudobulk distributions are bimodal compared to the normally distributed bulk data (Figure S3-4). This turns the problem more into a classification problem (is a gene low or high expressed), which might be easier to learn. On average, predicting cell population-specific expression is more difficult than predicting tissue-specific expression (Fig 1D-E): scXpresso_cp, pb p_erforms slightly worse than scXpresso_t, pb (_median correlation of 0.71 vs 0.75), but still better than scXpresso_t, b (_0.58).

One of the adaptations to Xpresso is that scXpresso_cp, pb i_s a multitask model. This slightly increases the performance compared to a single-task model (Figure S5) but mainly makes the model computationally more efficient. The marrow-FACS dataset, for instance, contains 22 cell populations. Since the single-task and multitask models need the same training time (approximately 30-60 minutes), this gives a 22x speed up.

The Tabula Muris scRNA-seq datasets were generated using two different protocols: 10X Genomics, a droplet-based method, and FACS-sorted Smart-seq2, a plate-based method. When comparing scXpresso_t, pb a_nd scXpresso_cp, pb t_rained on the two different protocols, e.g. lung-droplet vs. lung-FACS, we conclude that they perform equally well (Figure 1DE, S6-7). Depending on the tissue and cell population, one performs slightly higher than the other, but there are no significant differences. This is as expected since the pseudobulk values of both protocols are highly correlated (Pearson correlation > 0.85) (Figure S8). Hence, the protocol used to create the single-cell dataset does not influence the results.

For scXpresso_cp, pb,_ we tested how the two types of input features, DNA sequence and half-life time, influence the performance. We tested different lengths of the input sequence and whether one of the two features was enough to predict expression (Figure S9). A range of different sequence lengths results in the same performance (3.5-3.5, 7-3.5, and 10-5kb upstream-downstream). A longer sequence gives more information but also adds more noise. Since the model also becomes more complex, more parameters have to be learned and it takes more time and memory to train the model. Therefore, we decided to use 7kb upstream and 3.5kb downstream for further experiments. We also observed that adding the half-life time features results in higher performance, suggesting that these features are not easily captured from DNA sequences alone.

For the cell population-specific models, the performance varies considerably across different populations (Figure 1E). Comparing the populations in the lung dataset, for instance, the performance of the endothelial cells is very high compared to leukocytes (Figure 1F, S10). In general, the performance of scXpresso increases when more genes and cells are measured in a population (Figure 1F, S11). The leukocyte population is small (35 cells) and fewer genes are non-zero compared to other cell populations in the lung (8, 678 out of 20, 467 vs. 12, 715 on average). The ciliated cell population, on the other hand, is also small (25 cells), but this model reaches a higher performance. In this cell population, however, more genes were non-zero (11,717) compared to the leukocyte population. Hence, to train the model, we need a good representation of the cell population that includes enough expressed genes.

### Cell population-specific models outperform tissue-specific models

Now that we know that all models are well-trained, we predicted cell population-specific expression using the three different models to see whether increasing the resolution of the models increases the performance (Figure 2A). Since scXpresso_t, b a_nd scXpresso_t, pb w_ere trained using tissue-specific expression values, these models predict the same value for every cell population. On all datasets, scXpresso_cp, pb o_utperforms the tissue-specific models, which shows the benefit of training the models on a higher resolution (Figure 2B, S12A). Especially in more heterogeneous tissues, where the gene expression of cell populations is weakly correlated to the corresponding tissue, we see a large improvement (Figure 2C, S12B). For the lung-FACS dataset, for instance, the performance increases the most for immune cell populations (*Δ_cp_*_, *t*_ for B cells: 0.11, NK cells: 0.11, T cells: 0.09; see **Methods**) and the least for lung-specific populations (*Δ_cp_*_, *t*_ for stromal cells: 0.01, endothelial cells: 0.03, epithelial cells: 0.05). In the B cells in the lung, 4, 081 genes are not expressed in any of the cells and thus have a log-normalized expression of -4, but for which the tissue-specific model predicts a positive log-normalized expression value (Figure 2D). In contrast, the model trained on B cells predicts a lower expression for these genes (Figure 2E). Almost all these genes, however, are expressed in the lung (in the non-B cells), the lung-model learned this correctly too (Figure 2F).

**Figure 2.**
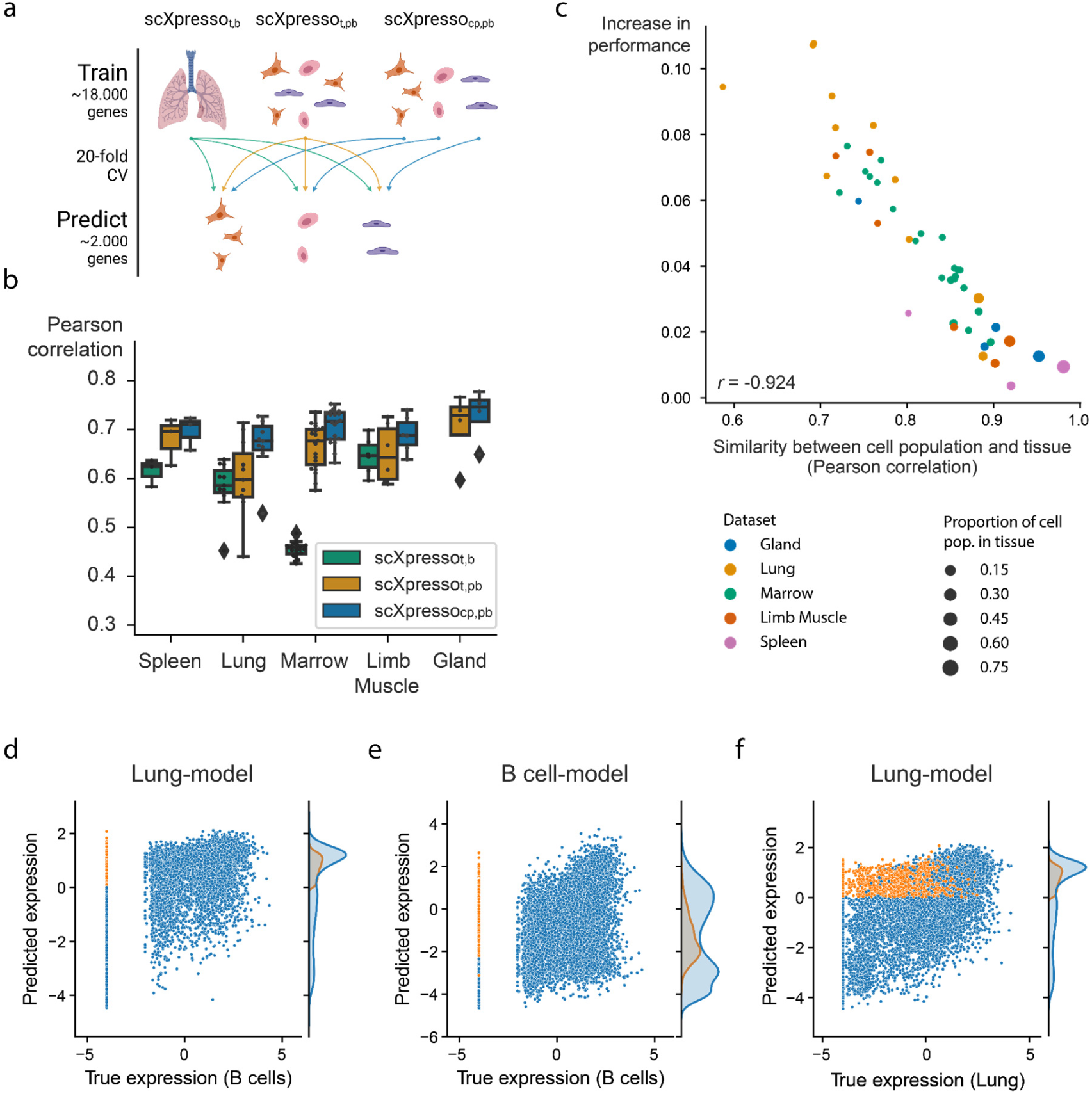
Comparison of the three scXpresso models for making cell population-specific predictions. **a)** Schematic overview of the experiment. **b)** Boxplot showing the performances of scXpresso_t, b (_tissue-specific (t) model on bulk (b) data), scXpresso_t, pb (_tissue-specific model on pseudo-bulk (pb) data), and scXpresso_cp, pb (_cell population-specific (cp) on pseudobulk (pb) data) on the cell population-specific task. Every point in the boxplot is the performance of a model on one cell population in that tissue (median Pearson correlation across the 20 folds). **c)** Similarity between a cell population and corresponding tissue (Pearson correlation between the true pseudobulk expression values) vs. the increase in performance (*Δ_cp_*_, *t*_, median Pearson correlation of scXpresso_cp, pb_ - scXpresso_t, pb_). Every dot is a different cell population and the colors represent the different tissues. **d-f)** Comparing the predictions made by the lung tissue model (lung-model) and the B cell population model (B cell-model). Genes where the lung-model predicts a too-high value are plotted in orange. **d-e)** True expression of the B cells vs. predicted expression by the **d)** lung-model and **e)** B cell-model. **f)** True expression of the lung cells vs. predicted expression of the lung model.

Some of the Tabula Muris datasets contain similar cell populations. For instance, B cells, macrophages, and T cells are measured in four, three, and three tissues, respectively. We hypothesized that if our models are cell population-specific, they should accurately predict the expression of a cell population in one tissue with a model trained on the same cell population but from another tissue (even though a cell’s tissue will slightly change the expression for (some) genes). To test this, we predicted the expression for common cell populations using three different types of models: 1) scXpresso_cp, pb t_rained on the same cell population, but from a different tissue, 2) scXpresso_cp, pb t_rained on a different cell population, but from the same tissue, 3) scXpresso_t, pb t_rained on the same tissue (Figure 3A). For example, we predict the expression of B-cells in the limb muscle, using 1) a model trained on B-cells in the lung, 2) a model trained on endothelial cells in the limb muscle, and 3) a model trained on the limb muscle. Again, the cell population-specific models outperform the tissue-specific models, even though they predict either a different dataset or a different cell population than they were trained on (Figure 3B, S13-14). This indicates that if you want to train a model for a cell population from a specific tissue where no single-cell data is available, you are better off using a model trained on a similar cell population from a different tissue than relying on a tissue-specific model. Whether a model trained on a different cell population and the same tissue performs better than a model trained on the same cell population but a different tissue, differs per tissue and cell population. For example, when predicting the expression of B cells in the limb muscle, the models trained on B cells in the marrow and lung even outperform the model trained on B cells in the limb muscle itself (Figure 3C). But, the models trained on different cell populations within the limb muscle perform variably when predicting B cells (Figure 3D). The models trained on immune populations, e.g. T cells or macrophages, perform similarly, but the muscle-specific populations perform worse. This difference between the B cell and the endothelial, mesenchymal stem cell, and skeletal muscle satellite cell models might seem small but is significant across the 20 folds (p-value = 9.5e-07 for all three populations, one-sided Wilcoxon rank sum test (26, 27)). Even though the differences are small, this indicates that our models indeed learn cell population-specific features.

**Figure 3.**
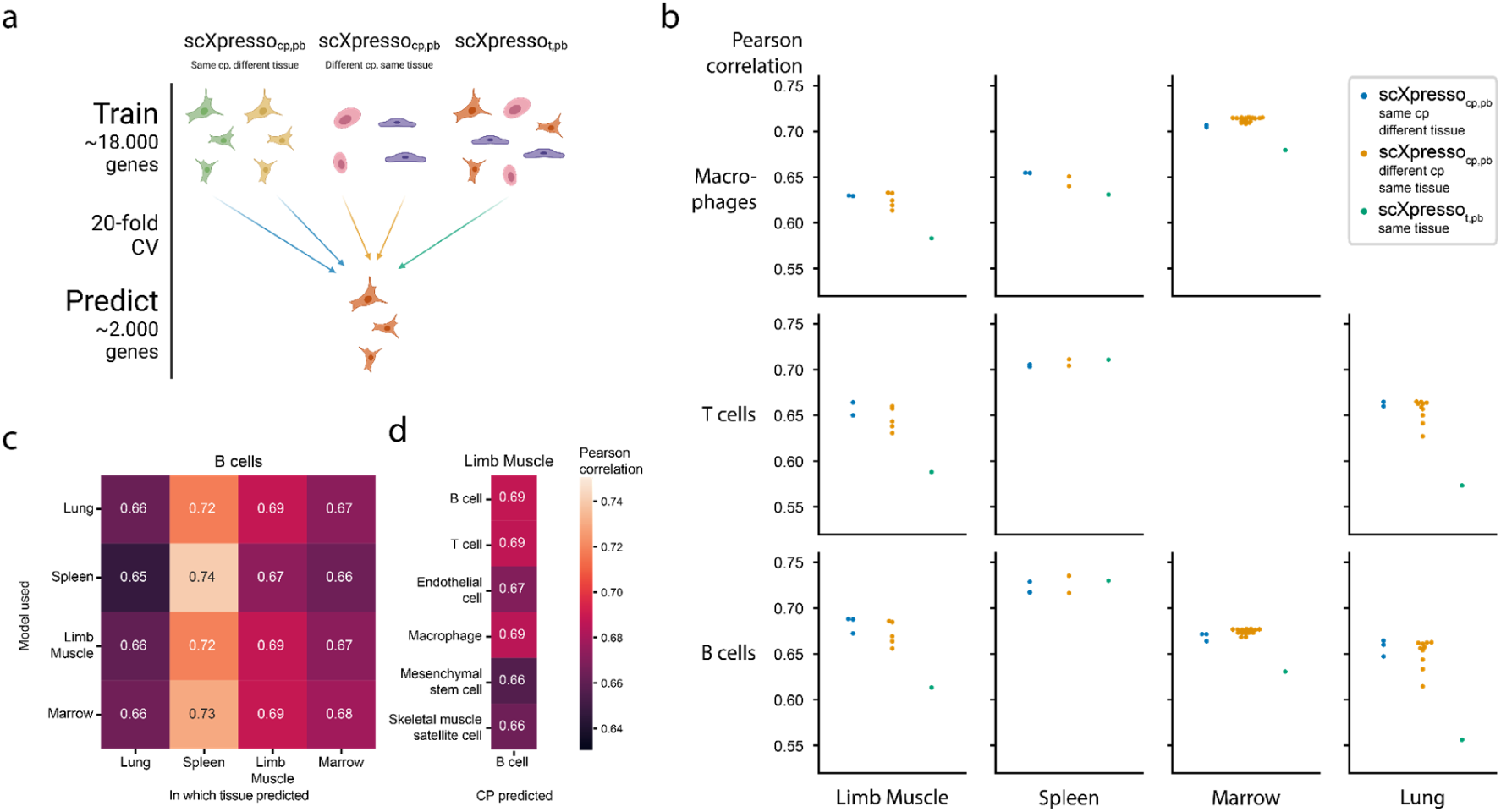
Comparing the predictions of scXpresso across cell populations and tissues. **a)** Schematic overview of the experiment. **b)** Performance (Pearson correlation) of three different types of models on different cell populations (rows) in different tissues (columns). Every dot is the median correlation of one model across the 20 folds. Since there are no T cells and macrophages defined in the Marrow and Lung dataset, these boxes are missing. **c)** Pearson correlation of different models when predicting the expression of B cells in different tissues. The rows indicate on which tissue scXpresso_cp, pb i_s trained, and the columns indicate for which tissue the expression of the B cells is predicted. **d)** Pearson correlation of different scXpresso_cp, pb w_hen predicting the expression of B cells in the limb muscle. Again the rows indicate which model is used.

### scXpresso learns expression patterns across human brain cell populations

Next, we applied scXpresso to a human brain dataset of the motor cortex (5). This dataset is annotated at different resolutions including a class (GABAergic, glutamatergic, and non-neuronal) and subclass (20 subclasses) level. Again, we trained models of different resolutions: a tissue- (t), class- (c), and subclass-specific (sc) model (scXpresso_t,_ scXpresso_c,_ and scXpresso_sc r_espectively). We used the trained models to predict the subclass-specific expression values (Figure 4A). Since scXpresso_t w_as trained on the tissue-specific pseudobulk expression, it predicts the same expression for all subclasses. The class-specific model, on the contrary, is a multitask model. Here, we use the predictions of the parent class to predict the expression of each subclass (i.e. subclasses belonging to the same parent class are predicted to have the same expression) (Figure S15). Similar to the Tabula Muris, we observed that increasing the resolution increases the performance: scXpresso_sc o_utperforms scXpresso_c w_hich outperforms scXpresso_t,_ (Figure 4B). For some subclasses, e.g. L2/3 IT, the performance barely improves when comparing scXpresso_sc w_ith scXpresso_c,_ which happens when the true expression values of the subclass and corresponding class are strongly correlated, similar as for the Tabula Muris case (Figure S16).

**Figure 4.**
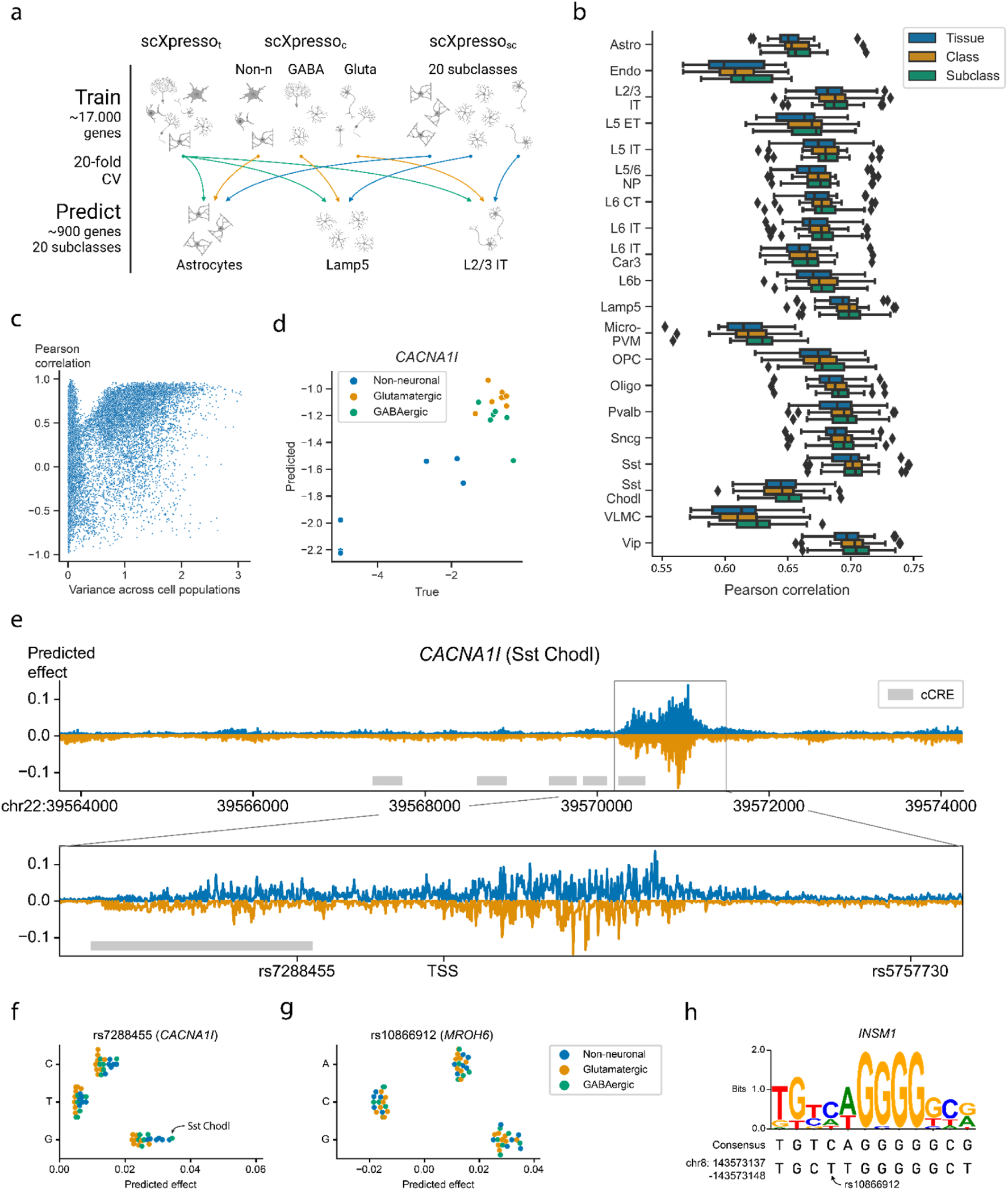
Performance of scXpresso on the human motor cortex. **a)** Schematic overview of the experiment. We train a tissue- (t), class- (c), and subclass-specific (sc) model (scXpresso_t,_ scXpresso_c,_ scXpresso_sc r_espectively) to predict the subclass-specific expression levels. **b)** Boxplots showing the Pearson correlation between the true and predicted values. Every point in the boxplot is the performance on a fold (n=20). **c)** Scatterplot showing the relation between the variance of a gene across the pseudobulk values of the subclasses and the Pearson correlation between the true and predicted values across the subclasses. Every dot is a gene. **d)** True and predicted expression for *CACNA1I.* Every dot is the expression in a subclass. Dots are colored according to their class. **e)** Mutation profile for *CACNA1I* for the Sst Chodl subclass. For every position, we calculated the difference in expression for all three possible substitutions and visualized the substitution with the highest absolute predicted effect. Mutations that are predicted to increase or decrease the expression are plotted in blue and orange, respectively. The grey rectangle highlights the region around the TSS. The grey boxes indicate the positions of candidate *cis*-Regulatory Elements (cCREs) derived from ENCODE data (25). **f-g)** Predicted effect, the predicted difference between the reference and alternative allele, of the three substitutions for **f)** rs7288455 on *CACNA1I* expression, and **g)** rs10866912 on *MROH6* expression. Every dot is one subclass and the dots are colored according to the class. **h)** Sequence logo and the consensus sequence for the *INSM1* transcription factor motif together with the sequence of the reference genome (bottom line).

Since genes with variable expression across subclasses are often interesting to study, we tested whether scXpresso_sc i_s able to learn the correct pattern for a gene across the subclasses. For every gene, we calculate the Pearson correlation between the true and predicted expression across the subclasses. If the expression of a gene shows some variance across the subclasses, scXpresso_sc p_redicts the pattern correctly (Figure 4C). An example is *CACNA1I*, a gene coding for a subtype of voltage-gated calcium channel that has been associated with schizophrenia (15, 28–31). Here scXpresso_sc c_orrectly learns that the expression in neuronal populations is higher than in non-neuronal (*r* = 0.90) (Figure 4D).

### *In-silico* mutagenesis reveals the most interesting GWAS variants

Since scXpresso can predict expression from the DNA sequence, we expect that it can also predict how the expression changes when the sequence is mutated. Therefore, we applied *in-silico* mutagenesis (ISM) to the sequence of *CACNA1I* and evaluated the predicted change in gene expression (6, 7, 11, 32). When comparing scXpresso_sc p_redictions for the Sst Chodl subclass across all possible mutations, we find mutations in the region around the TSS to affect the expression of the *CACNA1l* gene the most (Figure 4E). Note, that we did not input this location into the model, consequently, the model correctly identified that this is the most important region for transcriptional regulation. No other regions were found that affect the expression that strongly.

Besides visualizing the mutation pattern for one subclass, we can also visualize how ISM affects two subclasses differently. As an example, we compared the scXpresso_sc p_redictions for the Sst Chodl subclass and the L2/3 IT subclass (Figure S17). These predictions show that the Sst Chodl subclass is more sensitive to mutations than the L2/3 IT class for *CACNA1I*, which might be explained by the fact that *CACNA1I* is also higher expressed in Sst Chodl cells.

In addition, ISM can be used to prioritize variants of interest for diseases. *CACNA1I* is linked to 18 Schizophrenia-associated variants according to the NHGRI-EBI Catalog (33). Two of these variants, rs7288455 and rs5757730, fall within our input region (7kb upstream and 3.5kb downstream of the *CACNA1l* TSS). Mutating the reference A allele with the C or G variant at the position of rs7288455 increases the predicted expression for all cell populations (Figure 4F). The disease-associated variant, the A allele, is expected to decrease the expression (15, 33), which is in line with our predictions, although it is not known whether this is subclass-related. Our model suggests that the expression of *CACNA1I* increases the most in the Sst Chodl subclass. Interestingly, for the Sst Chodl subclass, this mutation results in one of the largest differences in *CACNA1l* expression amongst all other induced mutations (top 1% mutations with the strongest effect) (Figure S18). For the other variant, rs5757730, which lies in an intronic region, we see no difference in expression (Figure S19). Further supporting our predictions, rs7288455, but not rs5757730, overlaps with an ENCODE candidate *cis-*regulatory element. These results show that scXpresso can be used to prioritize GWAS hits.

In total, there are 3, 971 GWAS variants associated with Schizophrenia in the NHGRI-EBI Catalog (33). We focused on those genes that have two or more variants in the input region (20 genes, 49 variants) (Table S4). For these variants, we predicted the effect of all possible substitutions to prioritize the likely causal variants (Figure S20). For most genes, scXpresso predicts a profound effect for only one of the variants. For instance, when substituting ‘A’ with ‘C’ for the *HLA-B* variant rs2507989, the predicted expression of *HLA-B* decreases, while none of the mutations at the other variant positions of HLA-B, i.e. rs139099016 and rs1131275, are predicted to affect the expression. Noteworthy, rs1131275 is classified as a missense variant and thus not expected to alter transcription (33). For some genes, however, all variants seem to barely affect the expression.

Next, we checked if we could interpret the model predictions by characterizing the genomic sequences identified by scXpresso to have a strong effect on gene expression. For the *MROH-6* variant rs10866912, two substitutions are predicted to create an opposite effect. Substituting the reference ‘T’ with a ‘C’ is predicted to decrease the expression while mutating with a ‘G’ is predicted to increase the expression (Figure 4G). This variant is part of a binding site for the transcription factor *INSM1*, a transcriptional repressor (34) (Figure 4H). When substituting the ‘T’ with a ‘C’, the sequence of the reference genome becomes more similar to the consensus motif, while substituting with a ‘G’ makes the two sequences more dissimilar. This supports the predictions from scXpresso.

We compare our scXpresso predictions for these Schizophrenia variants to the predictions of Enformer, ExPecto, and Xpresso. For Enformer and ExPecto we used their pre-trained models which predict the expression for 5, 313 and 218 tissues/cell lines, respectively. Here, we only focused on the predictions related to the healthy brain (77 tracks for Enformer, 27 for ExPecto). For Xpresso, there were no pre-trained models for the brain available, so we trained the Xpresso model ourselves using bulk RNA-seq samples from the precentral gyrus, which is the region containing the motor cortex (see Methods). The expression values of the precentral gyrus are correlated to the pseudobulk expression values of the motor cortex (Figure S21A, r = 0.68). Similar to scXpresso, we used a 20-fold cross-validation to train the Xpresso model. The model is well-trained and reached a similar median correlation on the precentral gyrus as the scXpresso models on the motor cortex subclasses (Figure 4B, S21B-C, r = 0.69). Figure S22 shows the predictions for all models for the variants related to Schizophrenia. Using Xpresso and ExPecto we could not predict the effect of all variants, since some genes were missing from the data and some variants were lost during conversion from Hg38 to Hg19 (Table S4) (see Methods). It’s challenging to compare the predictions of the different methods since all models are trained on different brain regions or cell lines. Enformer usually predicts the same effect for the three different possible nucleotide mutations, e.g. for rs1131275 it predicts that all three substitutions decrease the expression. This variant, however, is classified as a missense variant, so we don’t expect it to alter transcription (33). For rs7288455, the variant close to *CACNA1I*, both scXpresso and Xpresso predict a similar effect, while Enformer and ExPecto predict only a very minimal effect. For rs10866912, the variant close to *MROH-6*, we showed that scXpresso could learn the TF binding site of *INSM1* while all the other models miss this pattern. These results overall illustrate the benefit of training prediction models on single-cell data.

## DISCUSSION

We presented scXpresso, a model to predict cell population-specific gene expression using the genomic sequence. We showed that scXpresso outperforms tissue-specific bulk and pseudobulk models especially when the expression profile of a cell population is dissimilar to that of the corresponding tissue. All scXpresso models reach a Pearson correlation of approximately 0.7 regardless of the cell population or tissue trained on. Additionally, the model learned the importance of the region around the TSS, transcription factor binding motifs (such as for *INSM1*), and the expression pattern of genes across different cell populations. Together, our findings show the potential of using single-cell data for predicting gene expression from sequence information in complex heterogeneous tissues.

We showed that it is possible to prioritize GWAS variants using scXpresso. Considering the expression of *CACNA1I*, we noticed that one variant, which overlaps with an ENCODE *cis-* regulatory element, is predicted to have a large effect, while another variant was predicted to have a negligible effect. The latter could be because the variant might affect splicing (which our model does not differentiate), the variant could be in a linkage disequilibrium block with other (associating) variants, or the variant could affect a more distant gene.

Comparing the predicted effects for mutations by scXpresso to other sequence-to-expression prediction models quantitatively is difficult as the true effect of these variants on specific brain regions and/or cell populations is unknown. We have shown that for a previously identified variant close to *CACNA1I* gene, both Xpresso and scXpresso predict an increase in expression, while ExPecto and Enformer predict a marginal effect. Note that, ExPecto and Enformer are not trained on specific brain regions, or cell population-specific data, but contain bigger structures such as the frontal cortex or frontal lobe. Hence, these models miss the cell population-specific effect of this variant. Hence, training these models on cell population-specific scRNA data could be an interesting next step.

Using our model, it is not possible to test trans-effects of variants as our model uses a limited genomic sequence region as input. Consequently, we could only test two variants related to Schizophrenia for *CACNA1I,* out of the 18 variants associated with *CACNA1I* (*33*). Ideally, we would increase the length of the input sequence, however, it is not easy to learn long-range interactions using CNNs. The Enformer model, which uses a 200kb sequence as input, tackles this problem by combining transformers and CNNs (11). Unfortunately, the Enformer model predicts CAGE reads instead of expression values, so we cannot trivially extend it or use it for single-cell data. An alternative approach might be to use their well-trained model to get an embedding for every input sequence and use this embedding to predict cell population-specific expressions.

Two future enhancements that we envision that could improve the performance of our model are related to the half-life time features and the output of the model. Currently, we extract five features from the mRNA sequence to approximate the half-life time. Recently, a new model, Saluki, was developed that could predict mRNA degradation rates directly from the sequence of the gene (38). Replacing the currently used features with those predicted by the Saluki model, or combining these features, might improve the cell population-specific predictions. A second potential improvement relates to the current output of scXpresso, which is the pseudobulk expression for every cell population, i.e. the average gene expression across all cells from that population. However, this ignores the variance within the population. It might be more beneficial to predict the distribution of gene expression across each population, instead of just one aggregated value.

In summary, we have shown the potential of predicting cell population-specific gene expression from genomic sequences by leveraging the resolution of single-cell data, opening the way for many new developments in this area.

## DATA AVAILABILITY

The pseudobulk expression values, trained models, and predictions are available on Zenodo: https://doi.org/10.5281/zenodo.7044908.

The code to reproduce the figures, train your own models, show the effect of variants, and do *in-silico* mutagenesis can be found on GitHub: https://github.com/lcmmichielsen/scXpresso.

## AUTHOR CONTRIBUTIONS

L.M., M.J.T.R., and A.M. conceived the study and designed the experiments. L.M. performed all the experiments and wrote the paper. L.M., M.J.T.R., and A.M. reviewed and approved the paper.

## ACKNOWLEDGEMENTS

We would like to thank Dr. Stavros Makrodimitris for his insightful discussion and his example of PyTorch code for convolutional neural networks. Figures 1B, 2A, 3A, 4A, and S15 were created with BioRender.com.

