## Supplement for "Predicting cell population-specific gene expression from genomic sequence"

### Supplementary figures

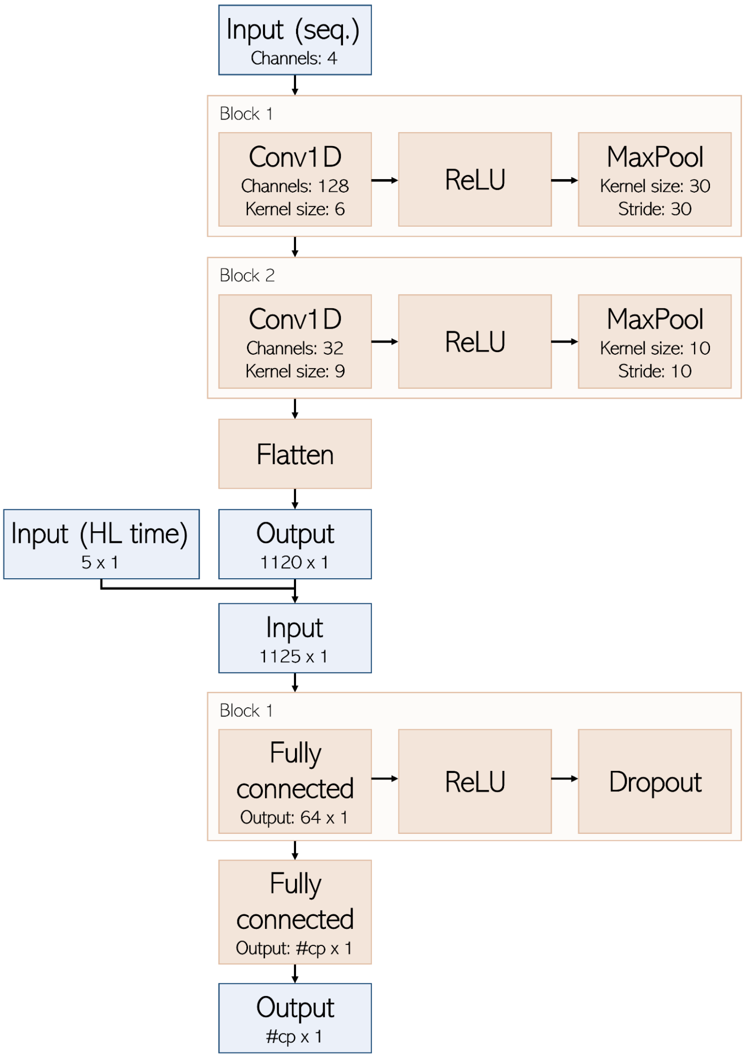


**Figure S1** Detailed overview of scXpresso. The orange blocks indicate the convolutional blocks and fully connected layers. The blue blocks indicate the input and output of the different parts.

a

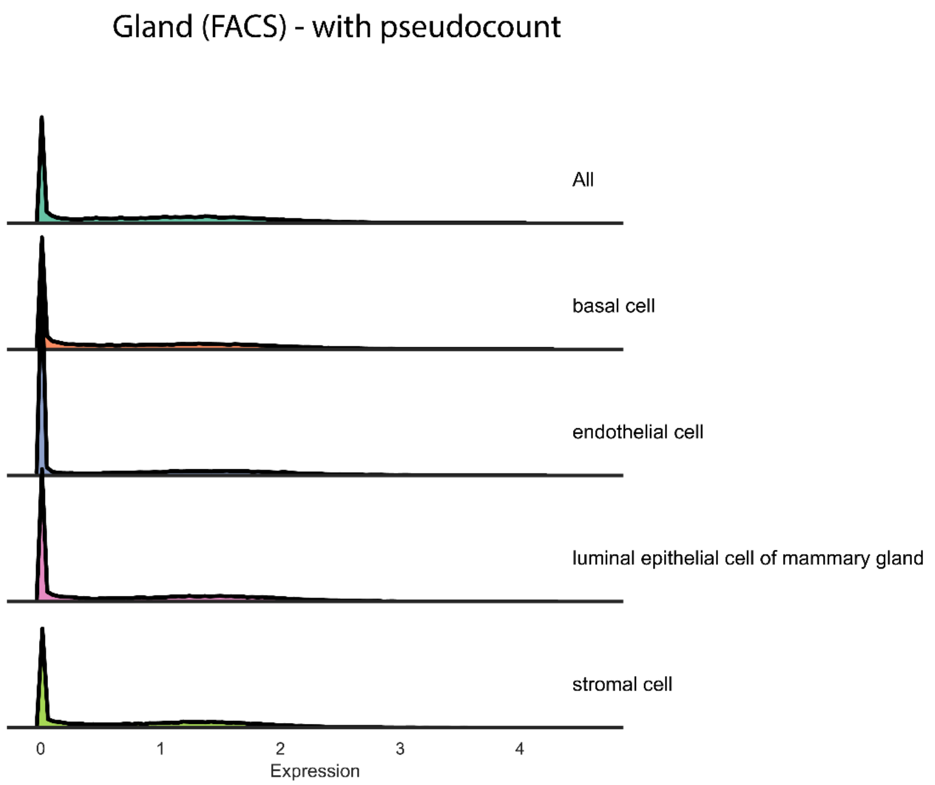


b

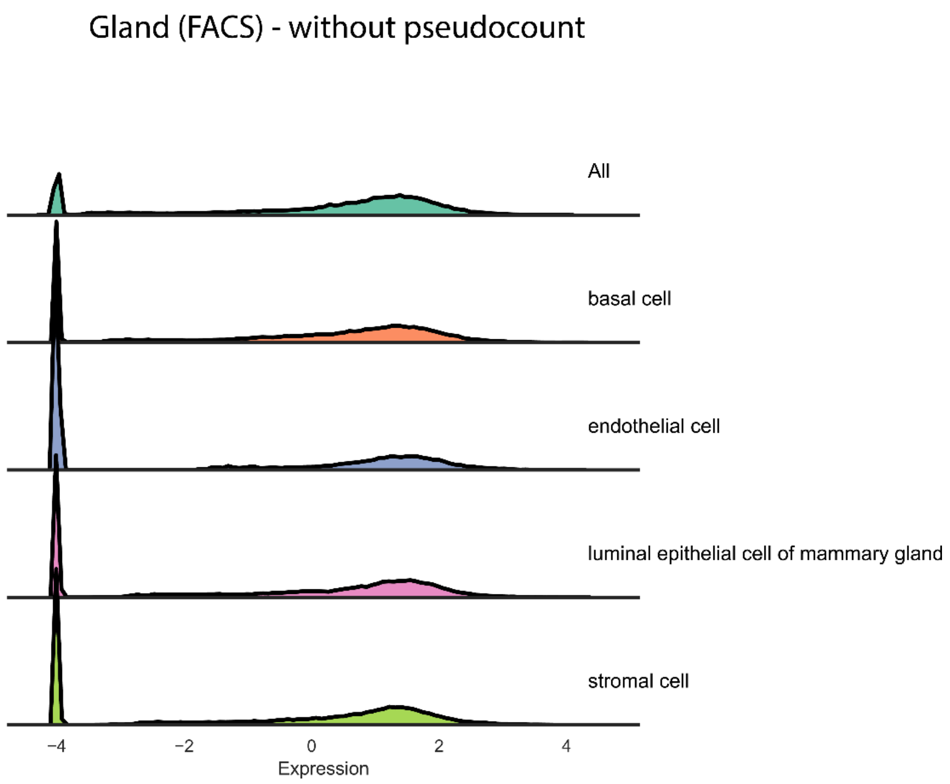


**Figure S2** Distribution of the pseudobulk expression when we **a)** add or **b)** do not add the pseudocount.

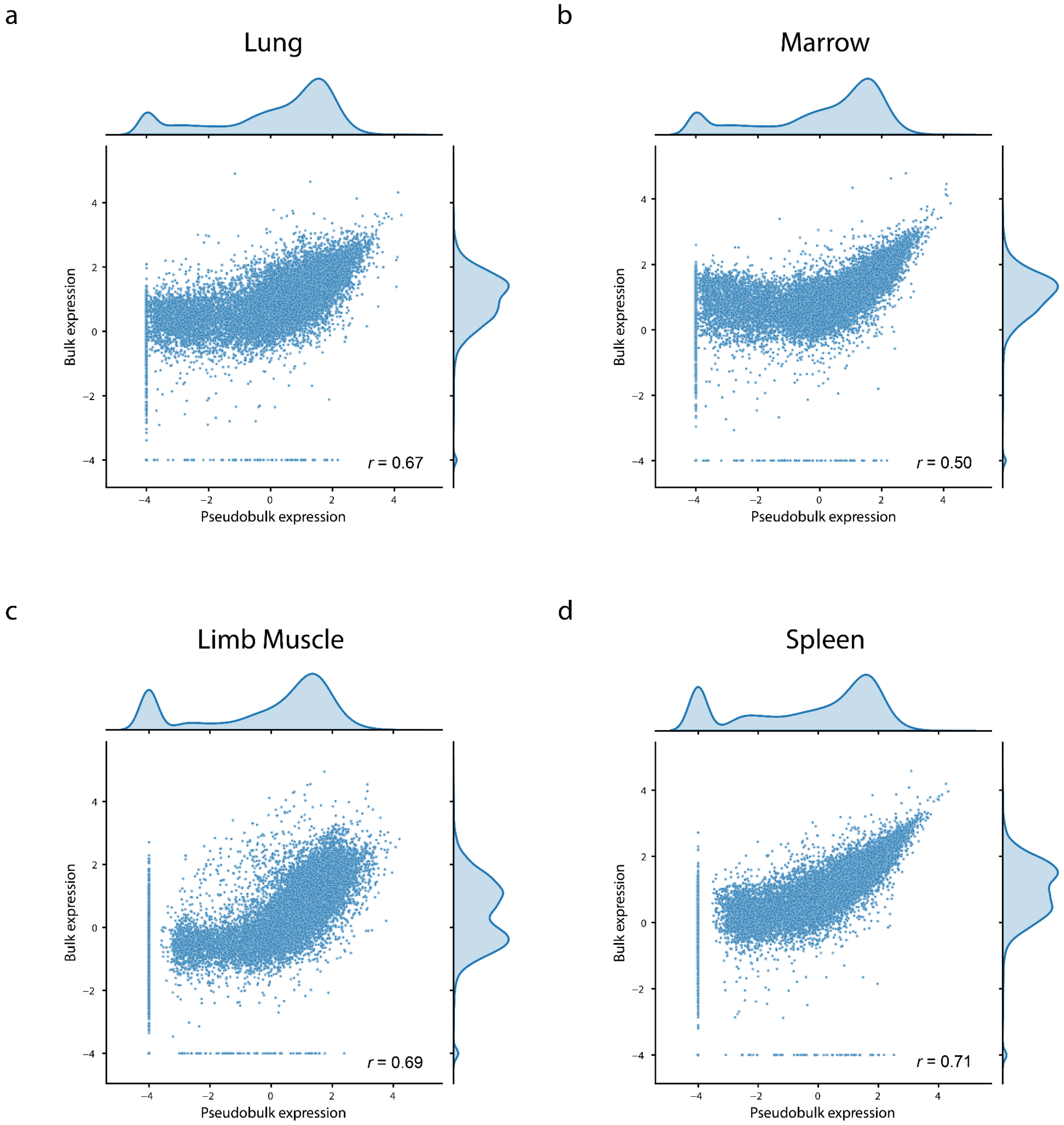


**Figure S3** Pearson correlation between single-cell (FACS-sorted Smart-seq2) pseudobulk and bulk expression values for the four different tissues: **a)** Lung, **b)** Marrow, **c)** Limb Muscle, and **d)** Spleen.

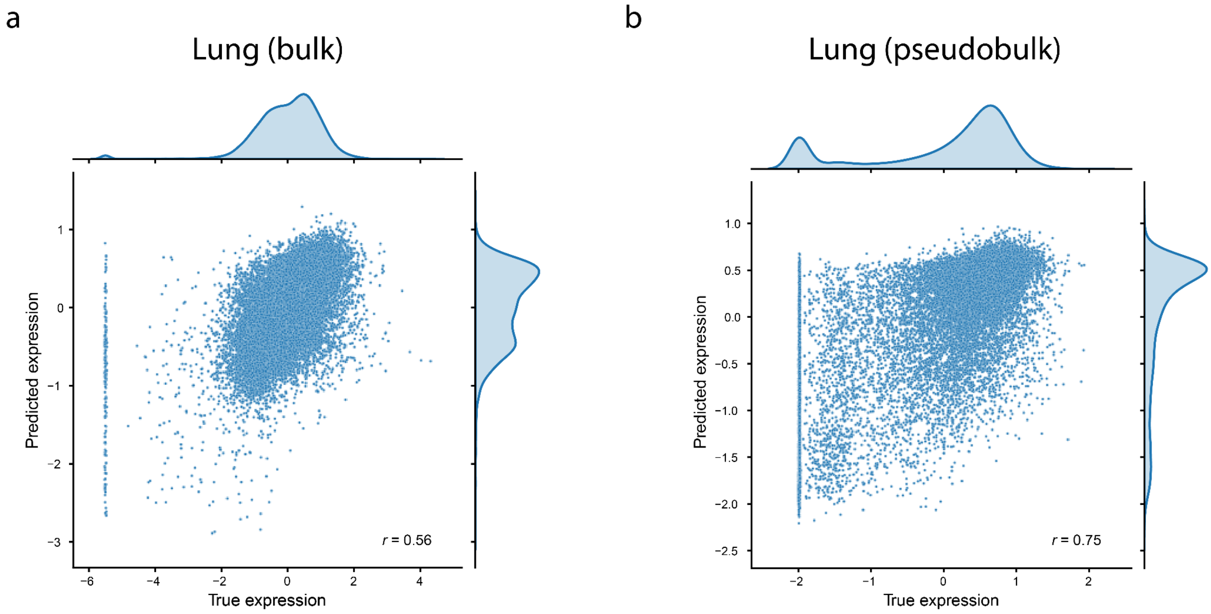


**Figure S4** Scatterplots showing the models' performance on individual tissues (lung). **a)** Performance of scXpresso$_{t,b}$ on the lung bulk dataset. **b)** Performance of scXpresso$_{t,pb}$ on the lung pseudobulk dataset.

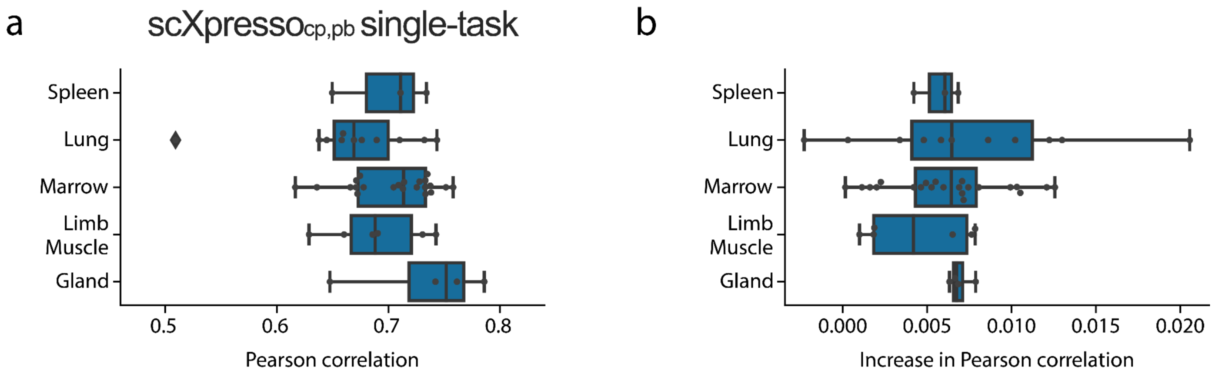


**Figure S5** Comparing the performance of the scXpresso$_{cp,pb}$ single-task, and multitask models. **a)** Pearson correlation of the scXpresso$_{cp,pb}$ single-task models. **b)** Increased performance when comparing the multitask model to the single-task models. Every dot represents the **a)** performance or **b)** increase in performance of the model on a cell population in that tissue (median Pearson correlation across the 20 folds).

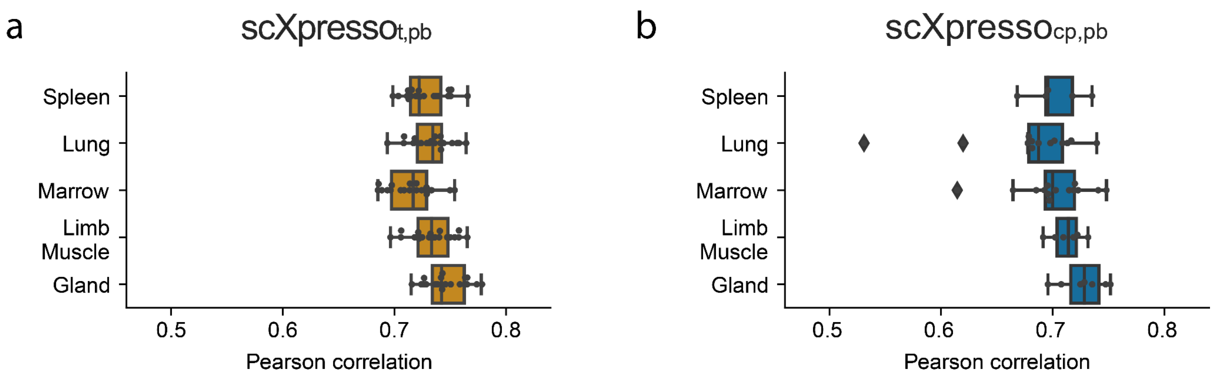


**Figure S6** Pearson correlation of **a)** scXpresso$_{t,pb}$, and **b)** scXpresso$_{cp,pb}$ on the droplet scRNA-seq datasets. For scXpresso$_{t,pb}$ every dot is the performance across one fold of the 20-fold CV. For scXpresso$_{cp,pb}$, we summarized the performance per tissue, so every dot represents the performance of the model on a cell population in that tissue (median Pearson correlation across the 20 folds).

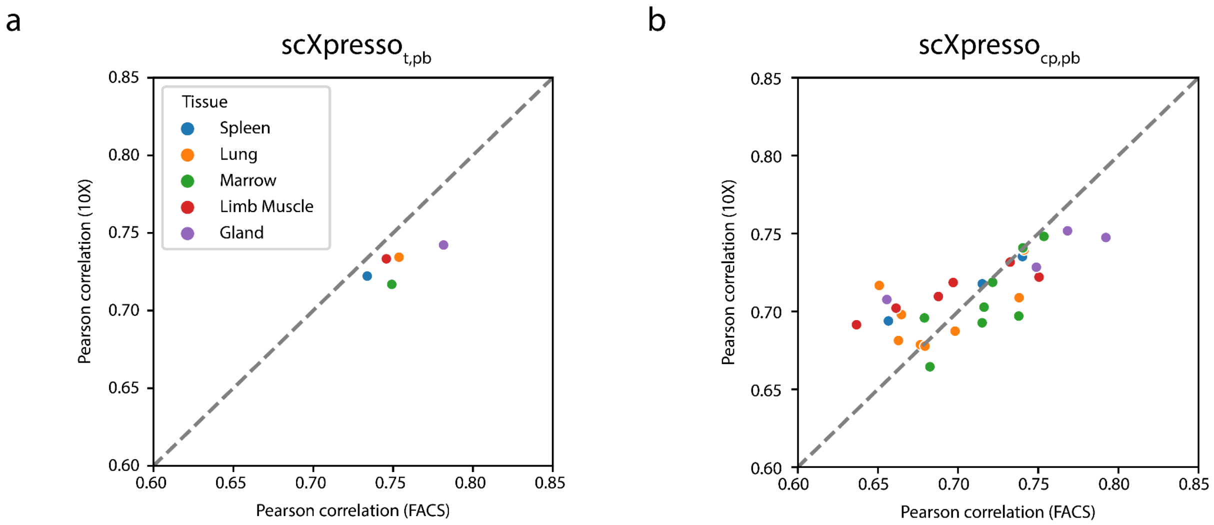


**Figure S7** Comparing the performance of **a)** scXpresso$_{t,pb}$ and **b)** scXpresso$_{cp,pb}$ on FACS-sorted Smart-seq2 and 10X Genomics data. Every point represents the median Pearson correlation across the 20 folds on a tissue (scXpresso$_{t,pb}$) or a cell population (scXpresso$_{cp,pb}$). The points' color represents the tissue of origin. In **b)** only the cell populations defined in both protocols are shown.

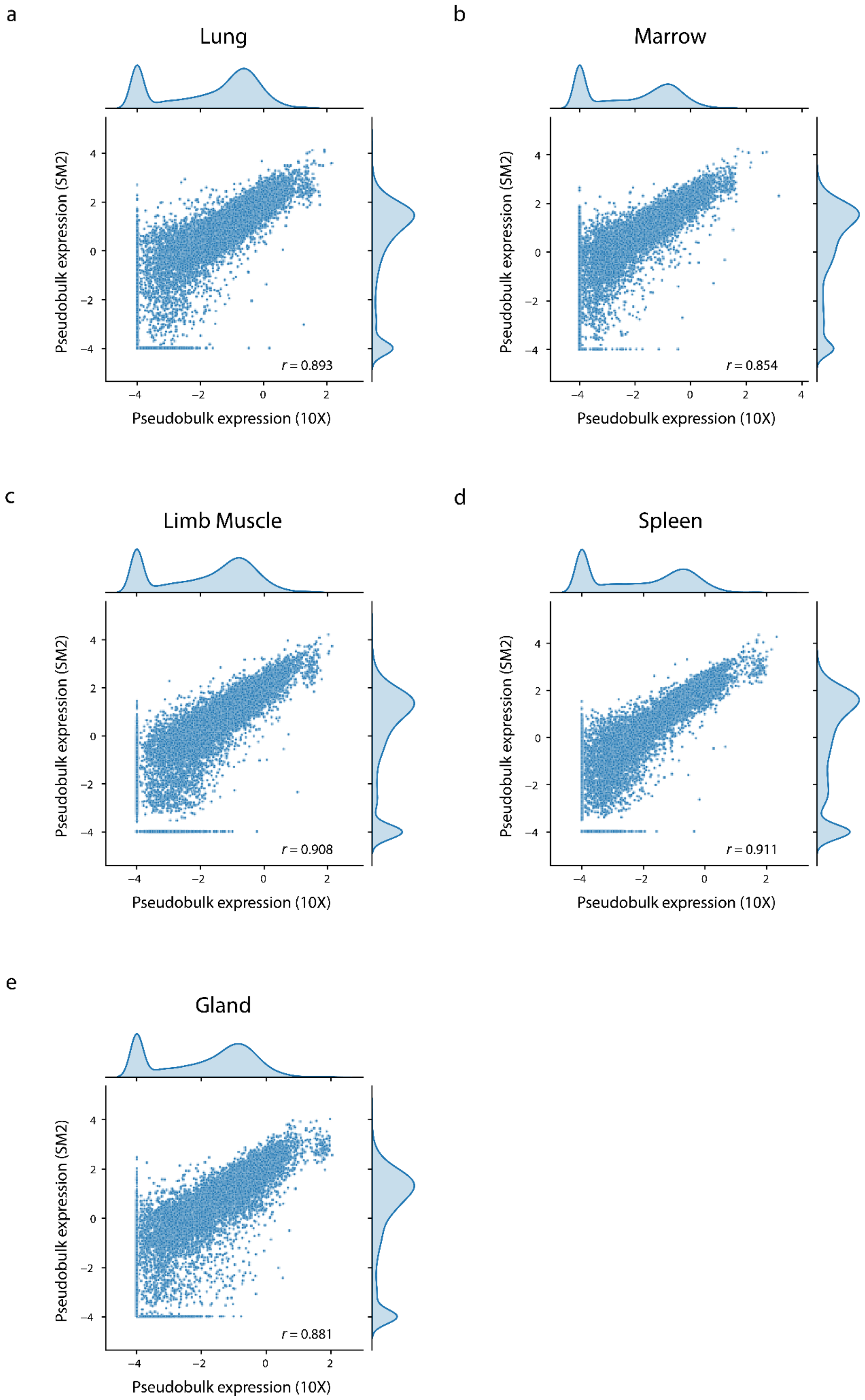


**Figure S8** Pearson correlation between 10X Genomics and FACS-sorted Smart-seq2 pseudobulk expression values for the five different tissues: **a)** Lung, **b)** Marrow, **c)** Limb Muscle, **d)** Spleen, and **e)** Gland.

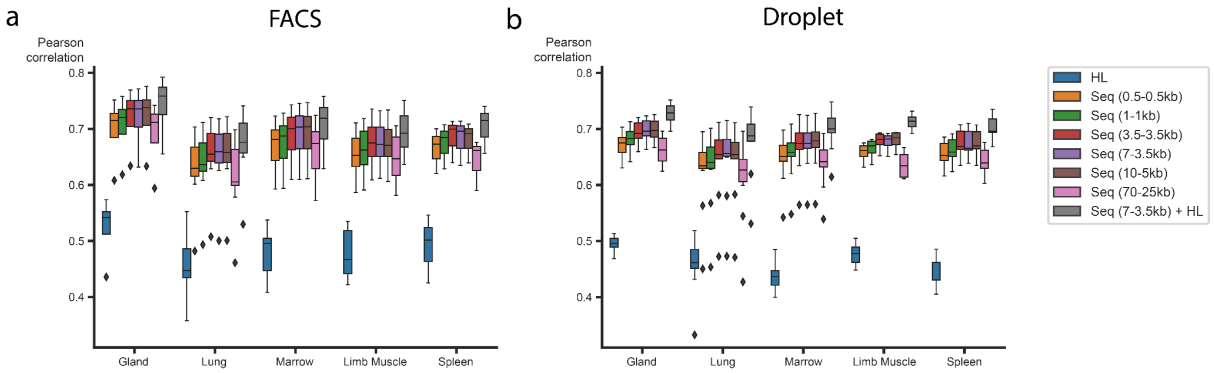


**Figure S9** Comparison of different input features for scXpresso$_{cp,pb}$. Performance of scXpresso$_{cp,pb}$ on the **a)** FACS Smart-seq 2 and **b)** 10X droplet datasets from the Tabula Muris. Points in the boxplot represent the model's performance on a cell population in that tissue (medium Pearson correlation across the 20 folds).

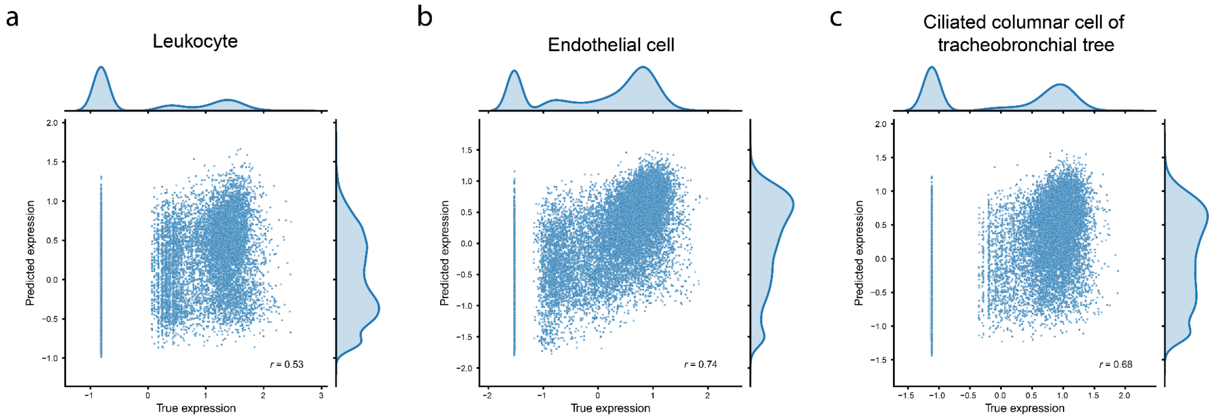


**Figure S10** Performance of scXpresso$_{cp,pb}$ on the **a)** leukocytes, **b)** lung endothelial cells, and **c)** ciliated columnar cells. The leukocytes & ciliated cells are both very small populations (35 and 25 cells respectively), but more genes are non-zero in the ciliated cells (8,678 and 11,717 non-zero genes respectively).

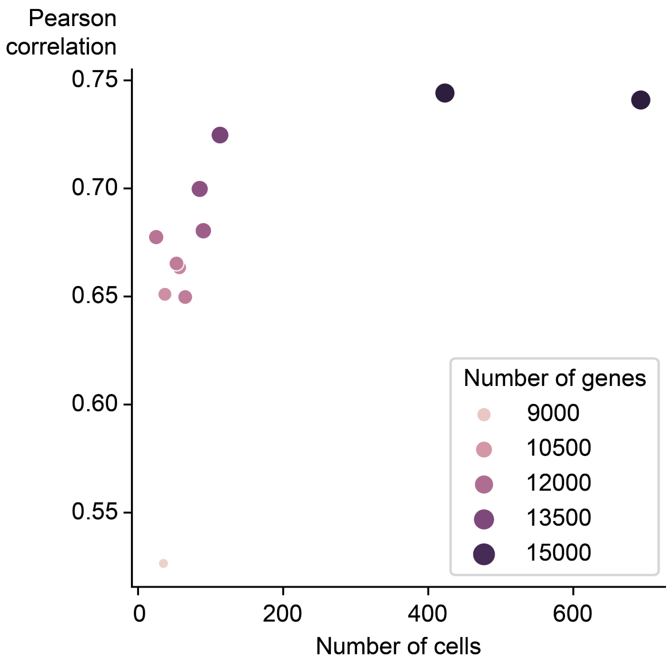


**Figure S11** Performance of scXpresso$_{cp,pb}$ on the cell populations in the lung dataset versus the number of cells per cell population. Every dot is the median Pearson correlation across the 20-folds on that cell population. The dots are colored and sized according to the number of non-zero genes in the cell population.

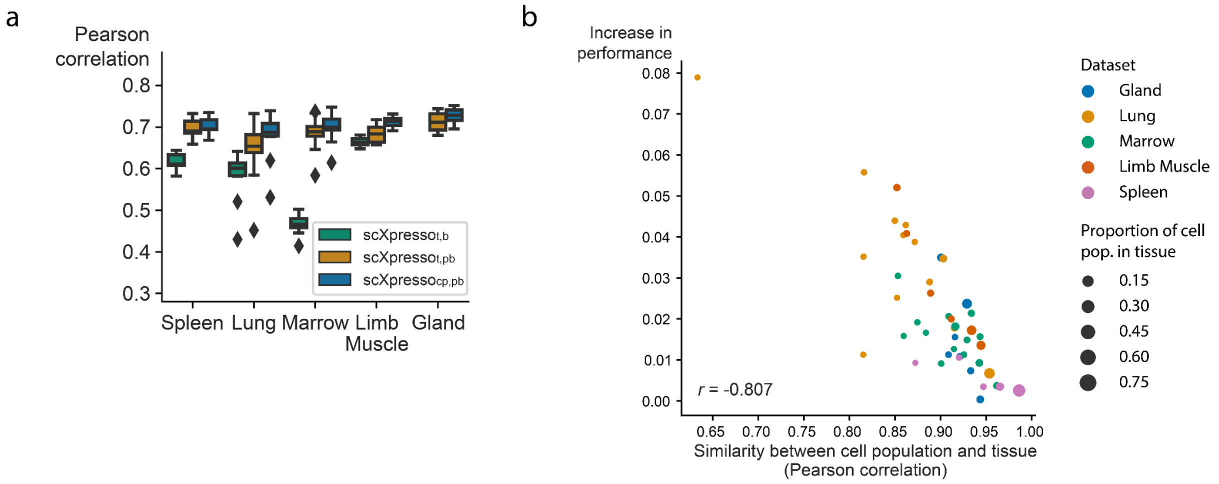


**Figure S12** Performance of scXpresso on the 10X droplet Tabula Muris datasets. Figure 2A explains the setup of the experiment. **a)** Boxplot showing the Pearson correlation of the three scXpresso models on the cell population-specific task. Every point in the boxplot is the model's performance on one cell population in that tissue (median Pearson correlation across the 20 folds). **b)** Similarity between a cell population and corresponding tissue (Pearson correlation between the pseudobulk expression values) vs. the increase in performance.

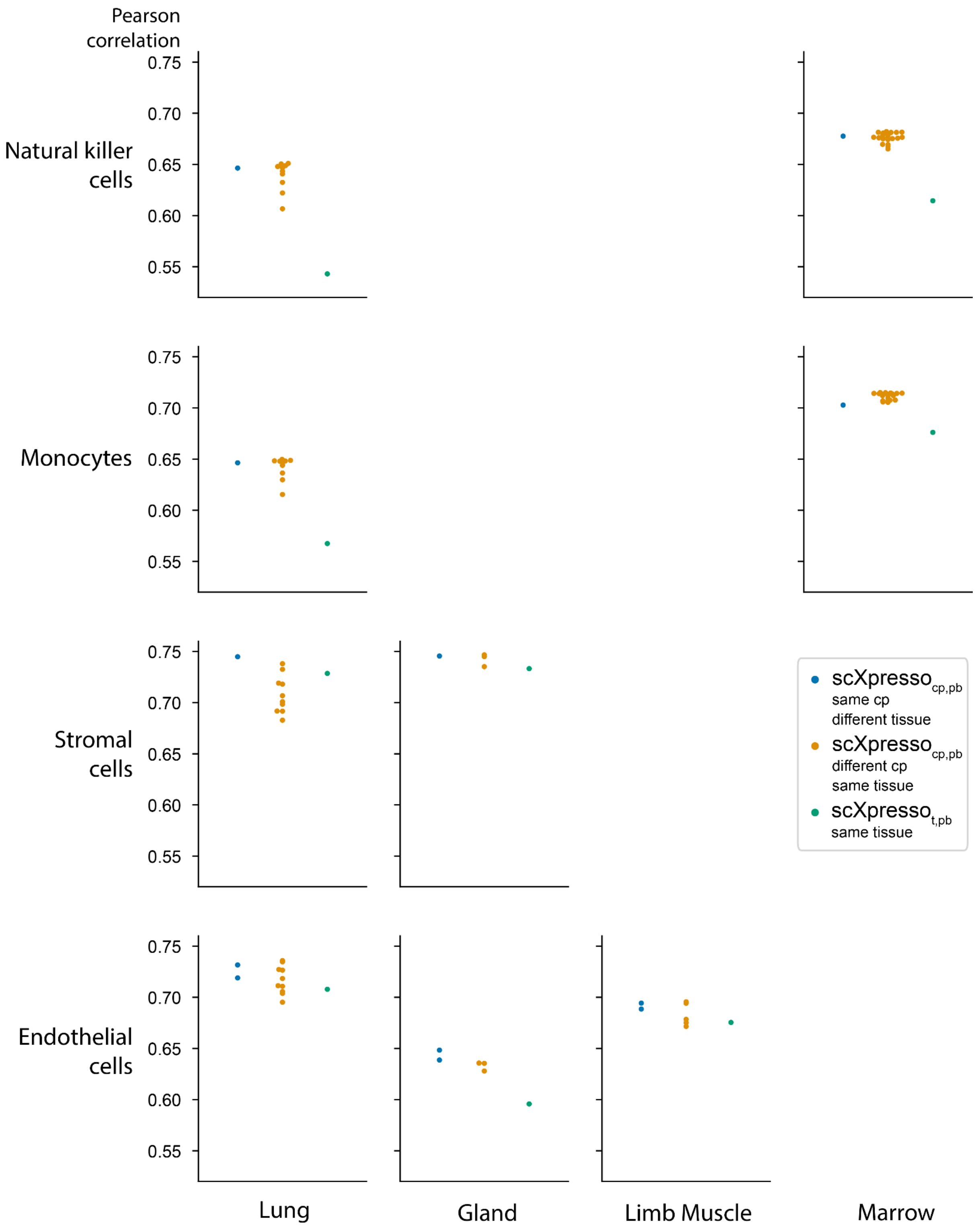


**Figure S13** Performance (Pearson correlation) of three different types of models on different cell populations (rows) in different tissues (columns). Every dot is the median correlation of one model across the 20 folds. The datasets used here are the Tabula Muris FACS-sorted datasets. Some panels are empty since not all cell populations are defined in all tissues.

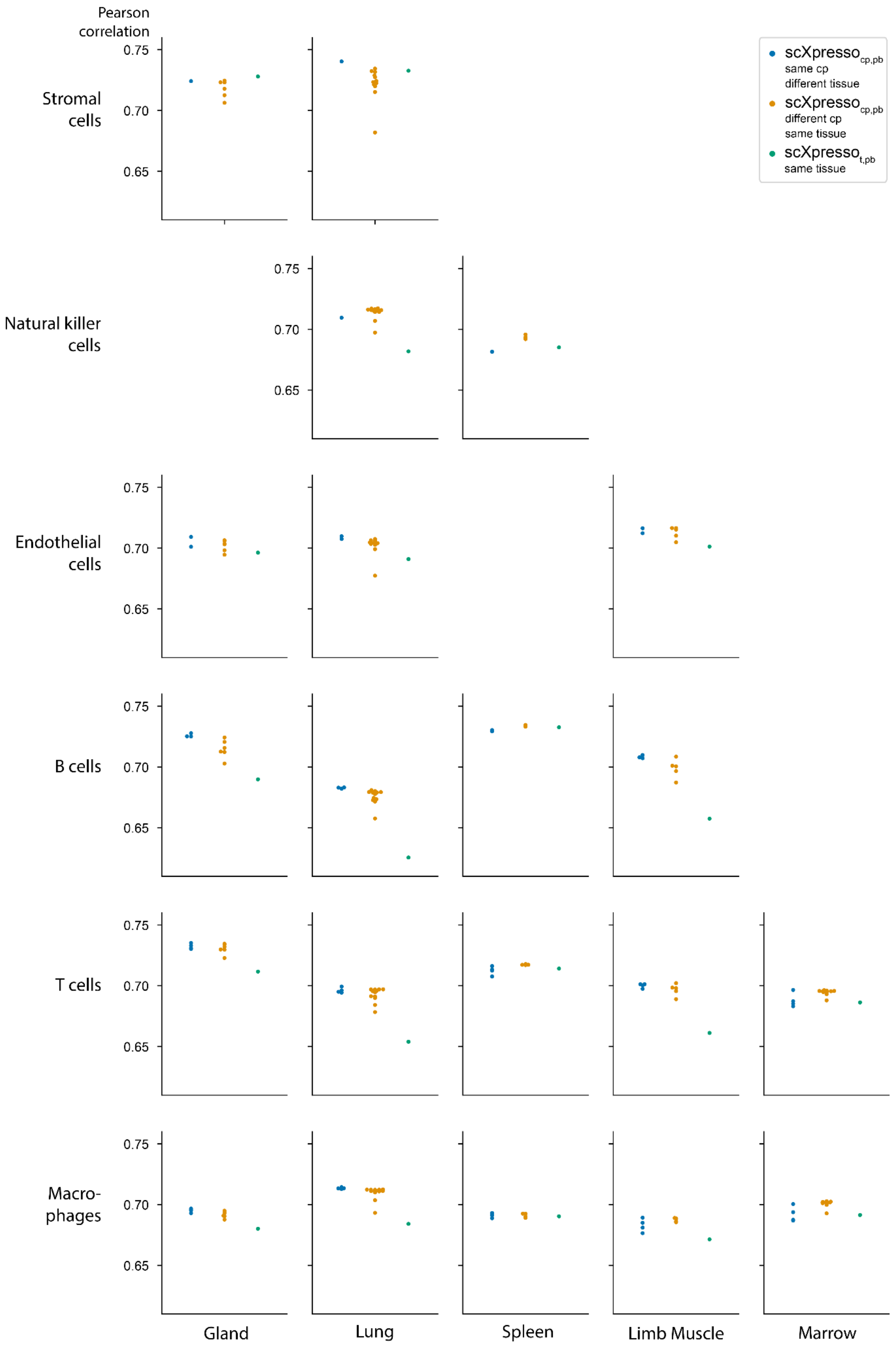


**Figure S14** Performance (Pearson correlation) of three different types of models on different cell populations (rows) in different tissues (columns). Every dot is the median correlation of one model across the 20 folds. The datasets used here are the Tabula Muris droplet datasets. Some panels are empty since not all cell populations are defined in all tissues.

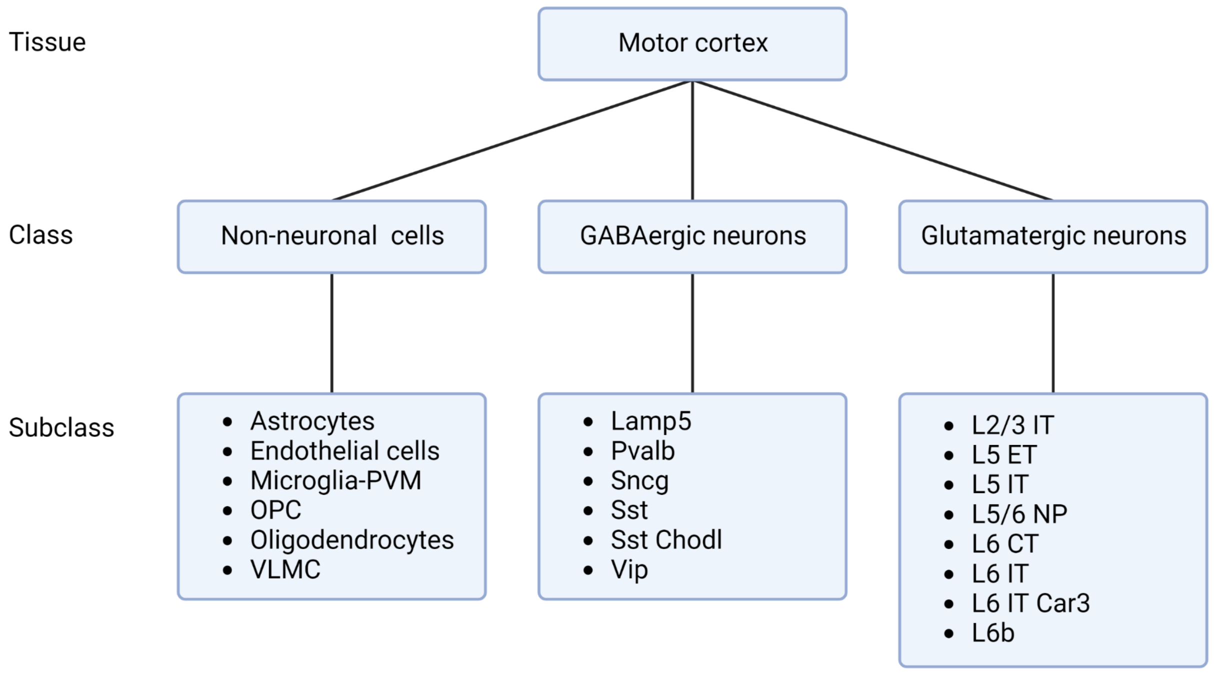


**Figure S15** Hierarchical representation of the classes and subclasses in the motor cortex.

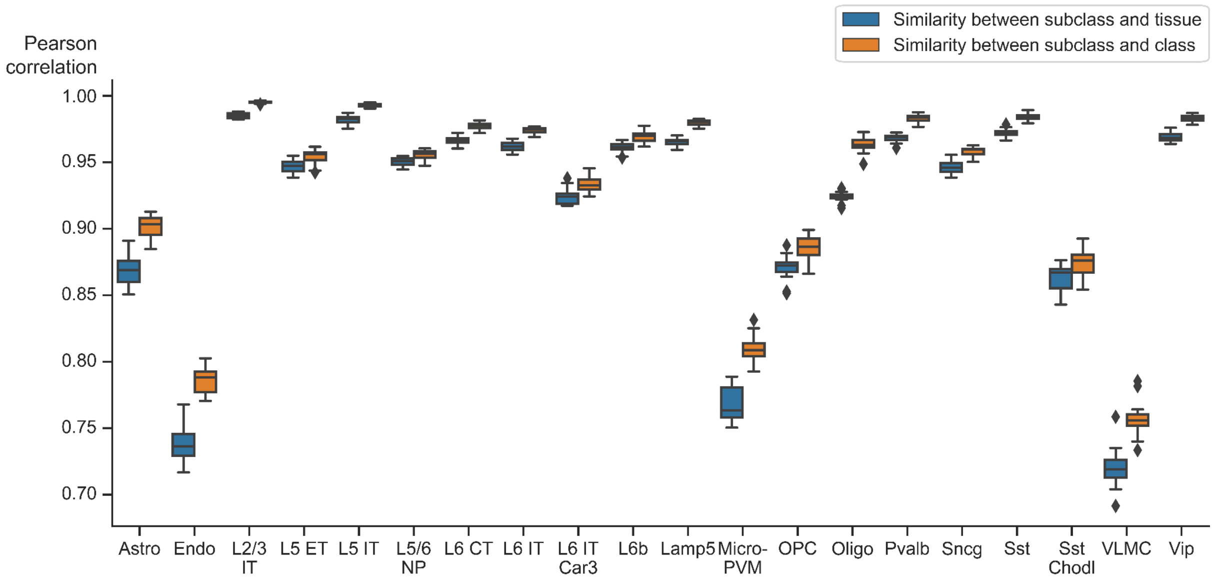


**Figure S16** Similarity between the true expression values of a subclass and the corresponding class and tissue (Pearson correlation). Every dot in the boxplot is the similarity across one fold of the 20-fold CV.

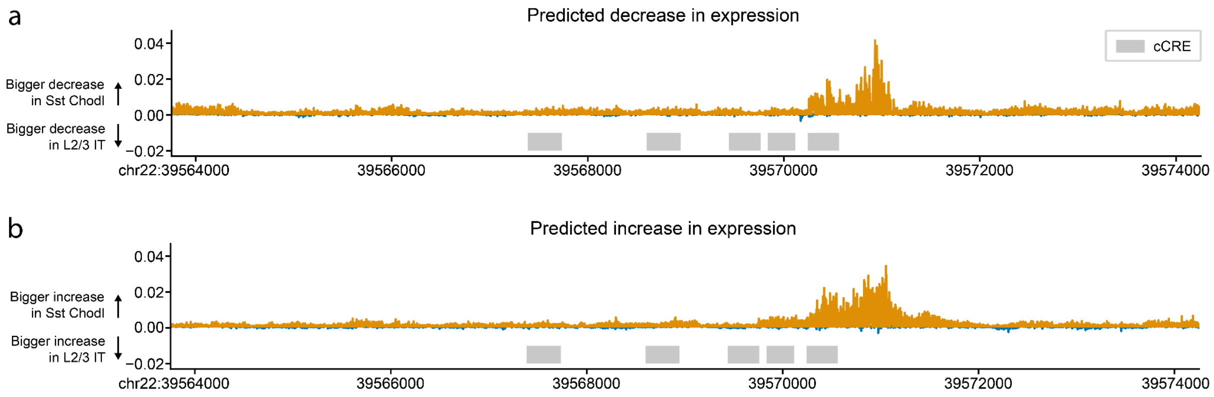


**Figure S17** Difference between the *in-silico* mutagenesis profiles of Sst Chodl and L2/3 IT subclass. In **a)** both subclasses are predicted to decrease in expression and in **b)** both to decrease in expression. Positive values (plotted in orange) mean that a bigger effect is predicted for Sst Chodl cells, while negative values (plotted in blue) mean that a bigger effect is predicted for L2/3 IT. The grey rectangles indicate the position of candidate cis-Regulatory Elements (cCREs) derived from ENCODE data.

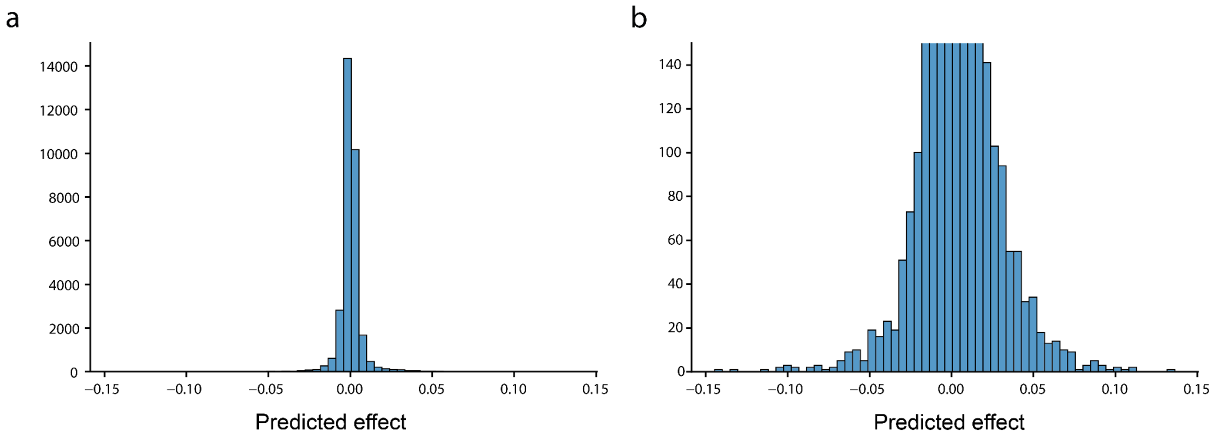


**Figure S18** Histograms showing **a)** the complete distribution and **b)** the focussed version of the predicted effect of all mutations (n = 31,500) for *CACNA1I* in the Sst Chodl subclass.

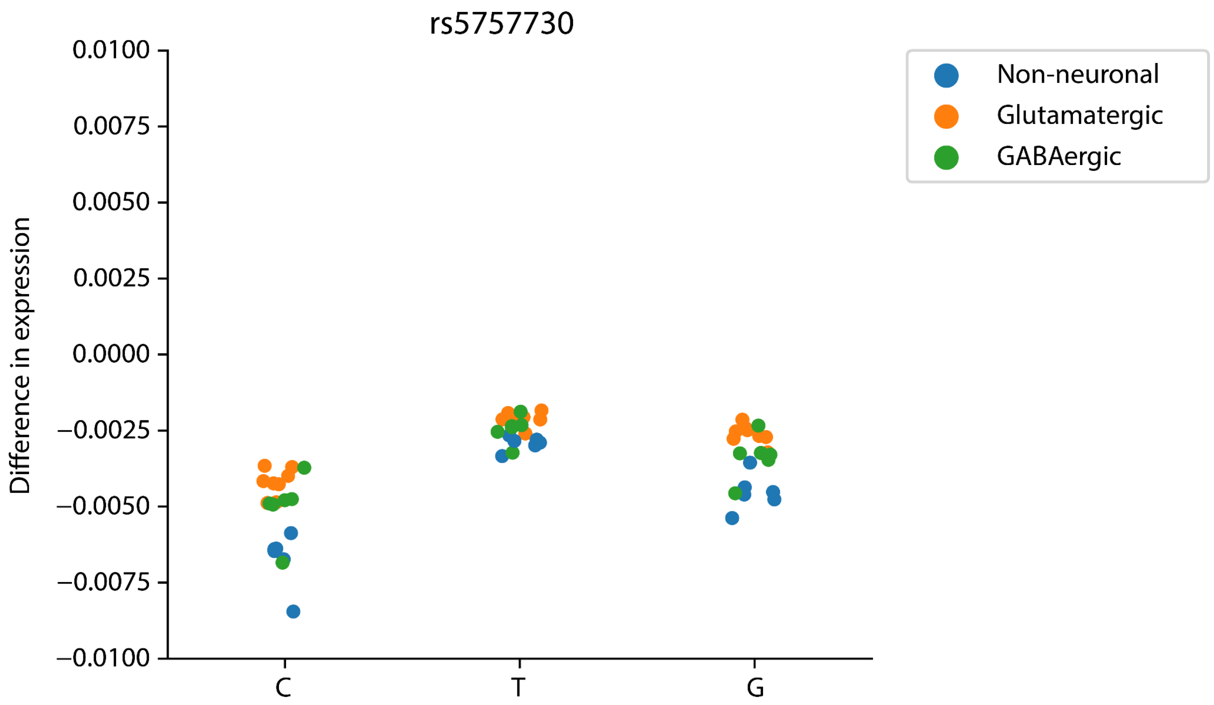


**Figure S19** The predicted effect of the three substitutions for rs5757730 on *CACNA1I* expression. Every dot is one subclass and the dots are colored according to the class. Note the scale on the y-axis. The predicted effect is very small.

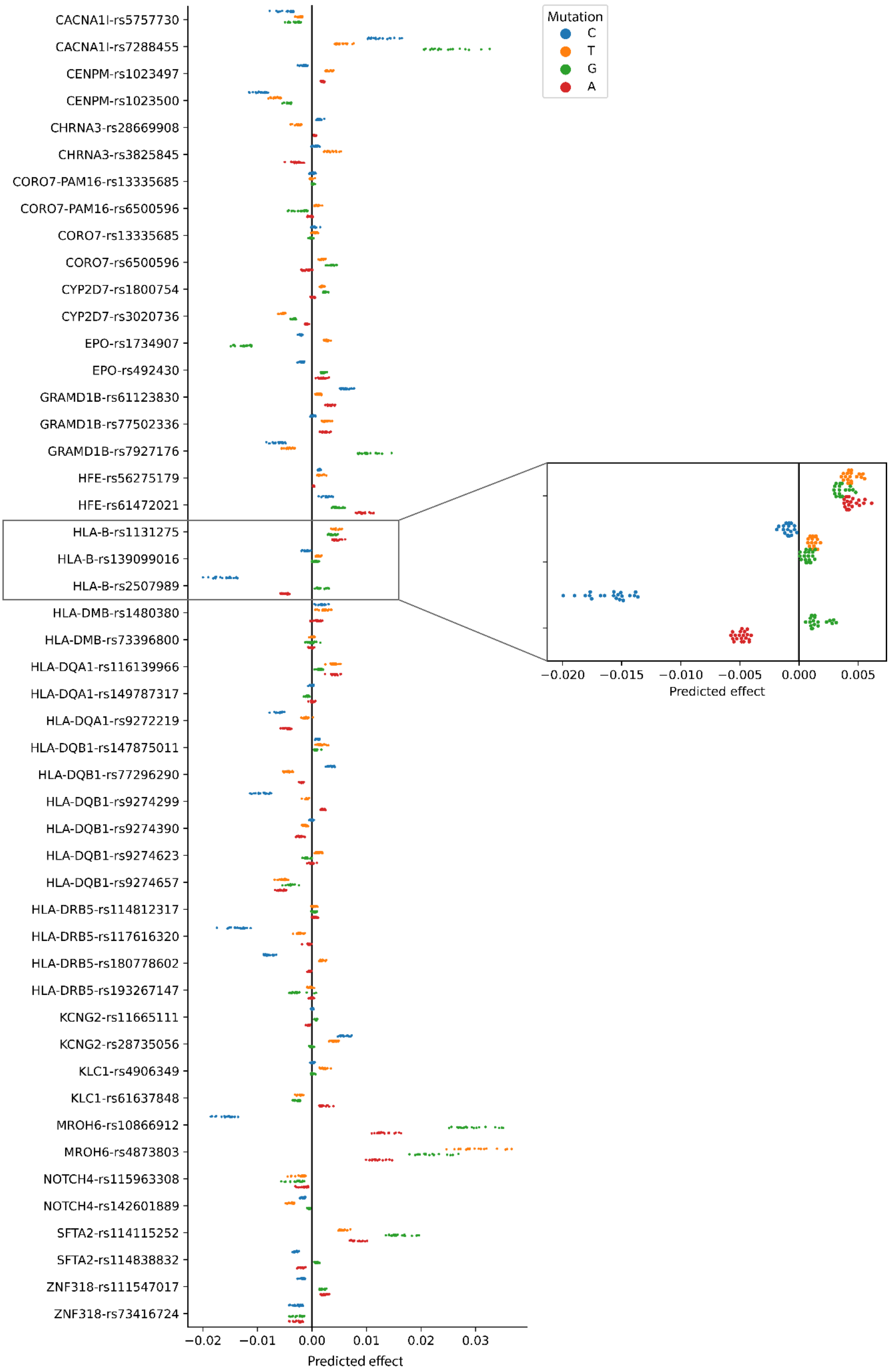


**Figure S20** We show the predicted effect for variants that are related to schizophrenia. For every variant, we indicate for which gene we predicted the effect. For every variant, you see three different substitutions (three colors). Every dot represents the predicted effect for one cell population.

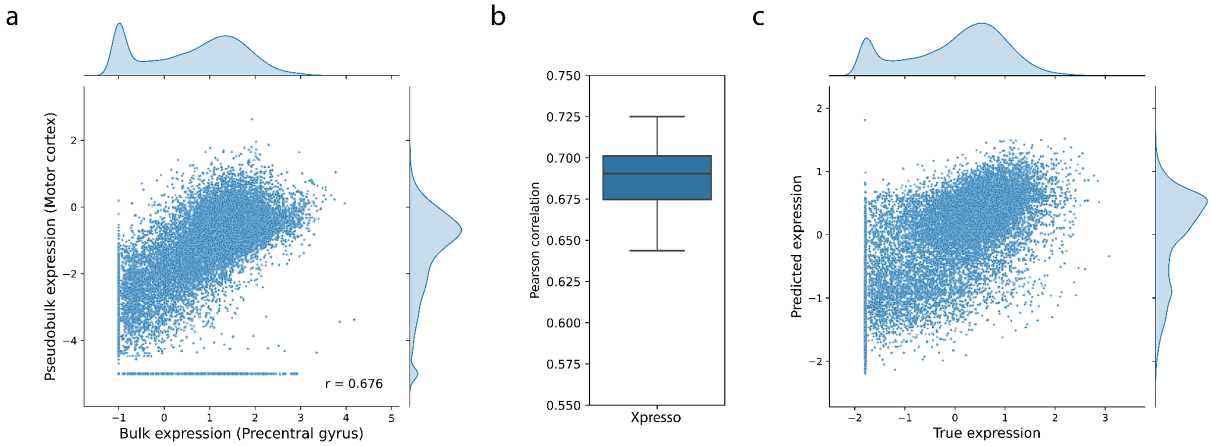


**Figure S21** Performance of Xpresso on the precentral gyrus dataset. **a)** Correlation between the bulk values for the precentral gyrus and the pseudobulk values for the motor cortex. **b)** Performance of Xpresso on the precentral gyrus dataset across 20-fold cross-validation. **c)** Scatterplot showing the predictions of Xpresso.

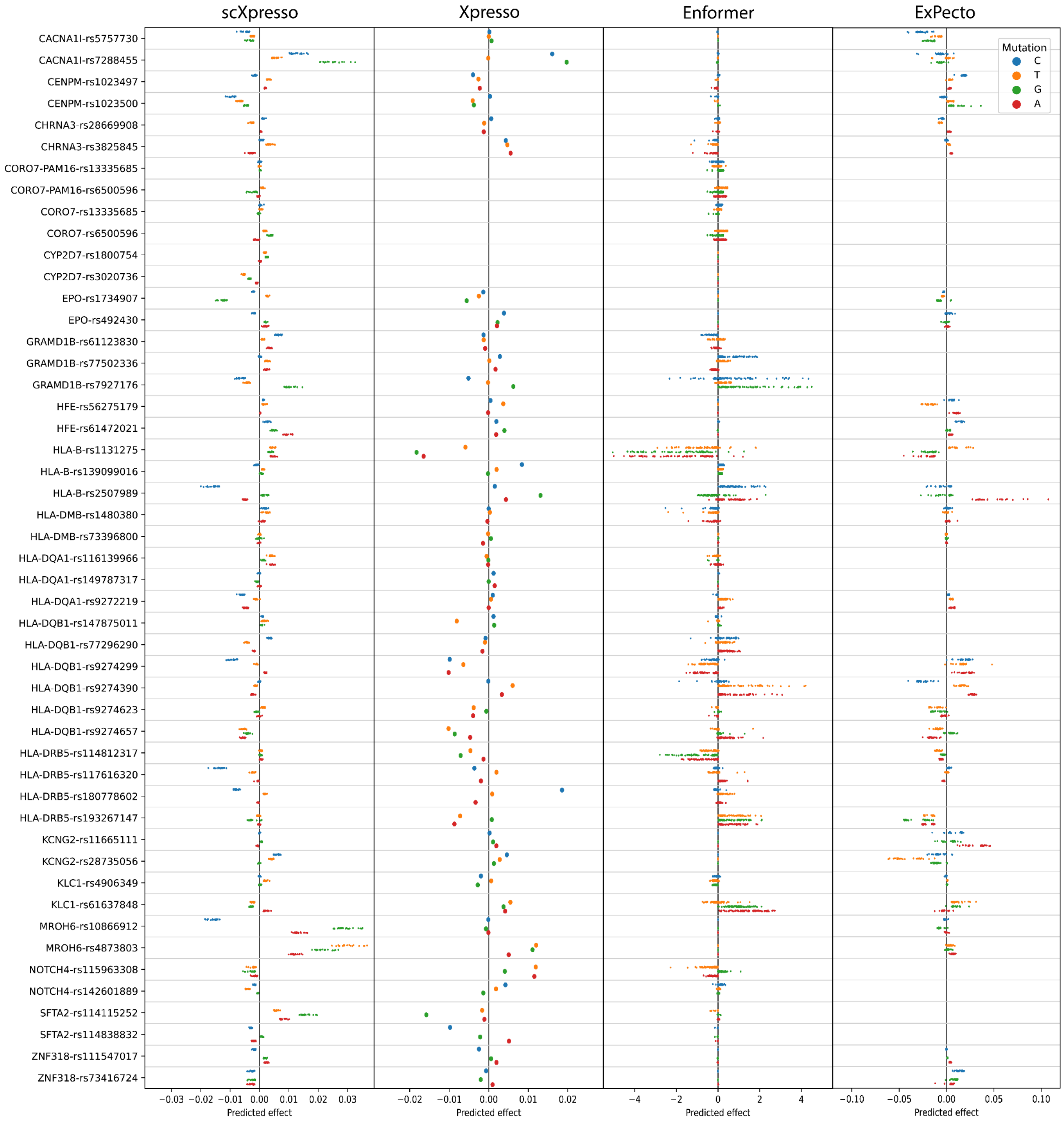


**Figure S22** The predicted effect for variants related to schizophrenia. For every variant, we predicted the effect using scXpresso, Xpresso, Enformer, and ExPecto. For scXpresso, every dot indicates the prediction of a subclass. For Enformer every dot indicates the prediction of a CAGE track related to the brain. For ExPecto every dot indicates the prediction on a certain brain tissue. For every variant, you see three different substitutions (three colors). Some variants are not predicted by all methods (see Table S3).

### Supplementary tables

**Table S1: Coding scheme for nucleotides**

| | **A** | **C** | **T** | **G** | **M** | **W** | **R** | **Y** | **S** | **K** | **B** | **D** | **V** | **G** | **N** |
|---|---|---|---|---|---|---|---|---|---|---|---|---|---|---|---|
| **A** | 1 | 0 | 0 | 0 | 1/2 | 1/2 | 1/2 | 0 | 0 | 0 | 0 | 1/3 | 1/3 | 1/3 | 1/4 |
| **C** | 0 | 1 | 0 | 0 | 1/2 | 0 | 0 | 1/2 | 1/2 | 0 | 1/3 | 0 | 1/3 | 1/3 | 1/4 |
| **T** | 0 | 0 | 1 | 0 | 0 | 1/2 | 0 | 1/2 | 0 | 1/2 | 1/3 | 1/3 | 0 | 1/3 | 1/4 |
| **G** | 0 | 0 | 0 | 1 | 0 | 0 | 1/2 | 0 | 1/2 | 1/2 | 1/3 | 1/3 | 1/3 | 0 | 1/4 |

**Table S2: List of CAGE tracks related to the brain in the Enformer model**

See Supplementary file.

**Table S3: List of ExPecto outputs related to the brain**

See Supplementary file.

**Table S4: List of variants related to Schizophrenia used**

See Supplementary file.

**Table S5: Overview of Tabula Muris datasets (single-cell) used during this study.**

| Tissue | Protocol | Genes (expressed) | Cells | Cell populations |
|---|---|---|---|---|
| Gland | 10X | 20467 (16039) | 4481 | 7 |
| | FACS | 20467 (17582) | 2663 | 4 |
| Limb Muscle | 10X | 20467 (15809) | 4543 | 6 |
| | FACS | 20467 (16742) | 1151 | 6 |
| Spleen | 10X | 20467 (14597) | 9573 | 5 |
| | FACS | 20467 (16725) | 1718 | 3 |
| Lung | 10X | 20467 (15954) | 5404 | 13 |
| | FACS | 20467 (17703) | 1923 | 11 |
| Bone marrow | 10X | 20467 (14682) | 4112 | 14 |
| | FACS | 20467 (18742) | 5355 | 22 |

**Table S6: Overview of Tabula Muris datasets (bulk) used during this study.**

| Tissue | Genes (expressed) | Samples |
|---|---|---|
| Limb Muscle | 20467 (20214) | 54 |
| Spleen | 20467 (20258) | 56 |
| Lung | 20467 (20282) | 55 |
| Bone marrow | 20467 (20263) | 54 |
